# Longitudinal single cell profiling of regulatory T cells identifies IL-33 as a driver of tumor immunosuppression

**DOI:** 10.1101/512905

**Authors:** Amy Li, Rebecca H. Herbst, David Canner, Jason M. Schenkel, Olivia C. Smith, Jonathan Y. Kim, Michelle Hillman, Arjun Bhutkar, Michael S. Cuoco, C. Garrett Rappazzo, Patricia Rogers, Celeste Dang, Orit Rozenblatt-Rosen, Le Cong, Michael Birnbaum, Aviv Regev, Tyler Jacks

**Author notes:** A.L., R.H.H., and D.C. contributed equally to this work.

## Abstract

Regulatory T cells (T_regs_) can impair anti-tumor immune responses and are associated with poor prognosis in multiple cancer types. T_regs_ in human tumors span diverse transcriptional states distinct from those of peripheral T_regs_, but their contribution to tumor development remains unknown. Here, we used single cell RNA-Seq to longitudinally profile conventional CD4^+^ T cells (T_conv_) and T_regs_ in a genetic mouse model of lung adenocarcinoma. Tissue-infiltrating and peripheral CD4^+^ T cells differed, highlighting divergent pathways of activation during tumorigenesis. Longitudinal shifts in T_reg_ heterogeneity suggested increased terminal differentiation and stabilization of an effector phenotype over time. In particular, effector T_regs_ had enhanced expression of the interleukin 33 receptor ST2. T_reg_-specific deletion of ST2 reduced effector T_regs_, increased infiltration of CD8^+^ T cells into tumors, and decreased tumor burden. Our study shows that ST2 plays a critical role in T_reg_-mediated immunosuppression in cancer, highlighting new potential paths for therapeutic intervention.

## INTRODUCTION

The recent clinical success of immune checkpoint inhibitors in the treatment of non-small cell lung cancer (NSCLC) highlights how targeting mechanisms of immunosuppression in the tumor microenvironment may be an effective therapeutic strategy (Makkouk and Weiner, 2015; Soria et al., 2015). However, only a subset of patients responds to immune therapies, suggesting that an improved understanding of other immunosuppressive mechanisms is needed for effective treatment.

One major mechanism of immunosuppression is posed by CD4^+^ regulatory T cells (T_regs_), which are thought to play a dominant role in impairing anti-tumor immune responses (Tanaka and Sakaguchi, 2017). T_regs_ are critical for maintaining peripheral immune tolerance and preventing autoimmunity (Josefowicz et al., 2012; Sakaguchi, 2011). Characterized by their expression of the transcription factor Foxp3, T_regs_ can inhibit adaptive immune responses through the production of inhibitory cytokines, direct killing of cells, competition with other T cell subsets for antigen or other substrates, and suppression of antigen presentation (Caridade et al., 2013; Savage et al., 2013; Vignali et al., 2008). T_regs_ are associated with poor prognosis in several cancers, including lung adenocarcinoma, which accounts for 40% of NSCLC (Fridman et al., 2012; Petersen et al., 2006; Shang et al., 2015; Shimizu et al., 2010; Suzuki et al., 2013). In mouse models, T_reg_ depletion can enhance anti-tumor immunity (Bos et al., 2013; Joshi et al., 2015; Marabelle et al., 2013), and antibodies directed against CTLA-4 act in part by depleting T_regs_ in the tumor microenvironment (Simpson et al., 2013).

While curbing T_reg_ function in tumors is an attractive therapeutic avenue, it is important to specifically target the tumor T_reg_ cell population to avoid systemic and potentially lethal autoimmune reactions. T_regs_ have considerable phenotypic diversity, which may help inhibit tumor-associated cells in different settings. Functional diversity within tumor T_reg_ populations may impact tumor immune responses, such that effector T_regs_ promote tumor growth (Green et al., 2017), whereas poorly immunosuppressive T_regs_ contribute to enhanced anti-tumor immunity (Overacre-Delgoffe et al., 2017; Saito et al., 2016). This functional diversity may be reflected in their transcriptional programs. Specific transcriptional profiles have been associated with T_regs_ in distinct tissues and inflammatory contexts, which are related to their tissue-resident functions (Arpaia et al., 2015; Burzyn et al., 2013; Cipolletta et al., 2012; Feuerer et al., 2009; Kolodin et al., 2015; Kuswanto et al., 2016). In human tumors, T_regs_ have a distinct program that may be shared across cancer types, and is associated with clinical outcome (De Simone et al., 2016; Magnuson et al., 2018; Plitas et al., 2016).

Inducible, autochthonous models of cancer are ideal for studying mechanisms of tumor tolerance because they recapitulate the longitudinal development of tumors and the immunosuppressive features of the endogenous tumor microenvironment better than transplanted, more “foreign”, tumors (Dranoff, 2011). Our group has previously developed a model of lung adenocarcinoma in which activation of oncogenic *K-ras*^*G12D*^ and loss of *Trp53* are driven by intratracheal delivery of a lentivirus expressing Cre recombinase (KP: *LSL-Kras*^*G12D*^, *p53*^*fl/fl*^) (DuPage et al., 2009; Jackson et al., 2005). By using lentivirus that also expresses firefly luciferase fused to chicken ovalbumin (Ova) and the antigenic peptide SIYRYYGL (Lenti-LucOS), we can program tumors to express known T cell antigens that can be used to monitor tumor-specific T cell responses (DuPage et al., 2011). Prior studies using this model have shown that T cell infiltration of Ova-expressing tumors delays tumor growth, but the number and activity of anti-tumor cytotoxic CD8^+^ T cells (CTLs) decline over time. The development of immune tolerance towards the tumor is partly due to the expansion of lung-resident T_regs_ that express various markers of effector activity and terminal differentiation (Joshi et al., 2015). T_reg_ depletion results in massive infiltration of CD4^+^ and CD8^+^ T cells into the lungs, suggesting that T_regs_ actively suppress anti-tumor immune responses. Since T_reg_-depleted animals succumb to systemic autoimmunity, a strategy targeting features of lung tumor-specific T_regs_ is required to minimize self-directed cytotoxicity.

Here, we map the phenotypic diversity of CD4^+^ T_conv_ and T_reg_ cells throughout tumor development in the KP model using scRNA-Seq. While T_conv_ subsets were stable over time, T_reg_ heterogeneity changed with tumor progression. At early time points, T_regs_ were less differentiated and expressed genes associated with interferon signaling, while mice with advanced disease had a greater proportion of effector T_regs_. Analyzing these data, we identified ST2 as a potential mediator of the accumulation of effector T_regs_ during tumor development. Indeed, T_reg_-specific ablation of ST2 increased CD8^+^ T cell infiltration of tumors and reduced tumor size while avoiding systemic autoimmunity. Our high-resolution characterization of T_reg_ heterogeneity in the tumor microenvironment thus allows us to define refined and effective ways to target T_reg_ function in cancer.

## RESULTS

### CD103 and KLRG1 mark an activated, heterogeneous population of lung tissue T_regs_

We have previously demonstrated that tumor development in the KP model is associated with the expansion of lung-infiltrating T_regs_, a large proportion of which express CD103 (integrin aE) and killer cell lectin-like receptor 1 (KLRG1), which have been associated with T_reg_ effector activity and terminal differentiation, respectively (Beyersdorf et al., 2007; Cheng et al., 2012; Huehn et al., 2004; Lehmann et al., 2002; Sather et al., 2007). We characterized the heterogeneity of the T_reg_ population in KP mice with advanced disease. While T_reg_ cells in the draining lymph node (dLN) were predominantly CD103^+^KLRG1^−^(double-negative, DN) or CD103^+^KLRG1^−^(single-positive, SP), nearly 40% of lung T_regs_ from late-stage, tumor-bearing KP mice were CD103^+^KLRG1^+^ (double-positive, DP) (**Figure 1A**). DP T_regs_ in late-stage, tumor-bearing mice had increased expression of genes associated with enhanced T_reg_ cell activity, including GITR, CD39, and PD-1, compared to SP and DN T_regs_ (Joshi et al., 2015). We therefore hypothesized that these T_reg_ subsets may have distinct tissue and tumor-specific transcriptional programs.

**Figure 1.**
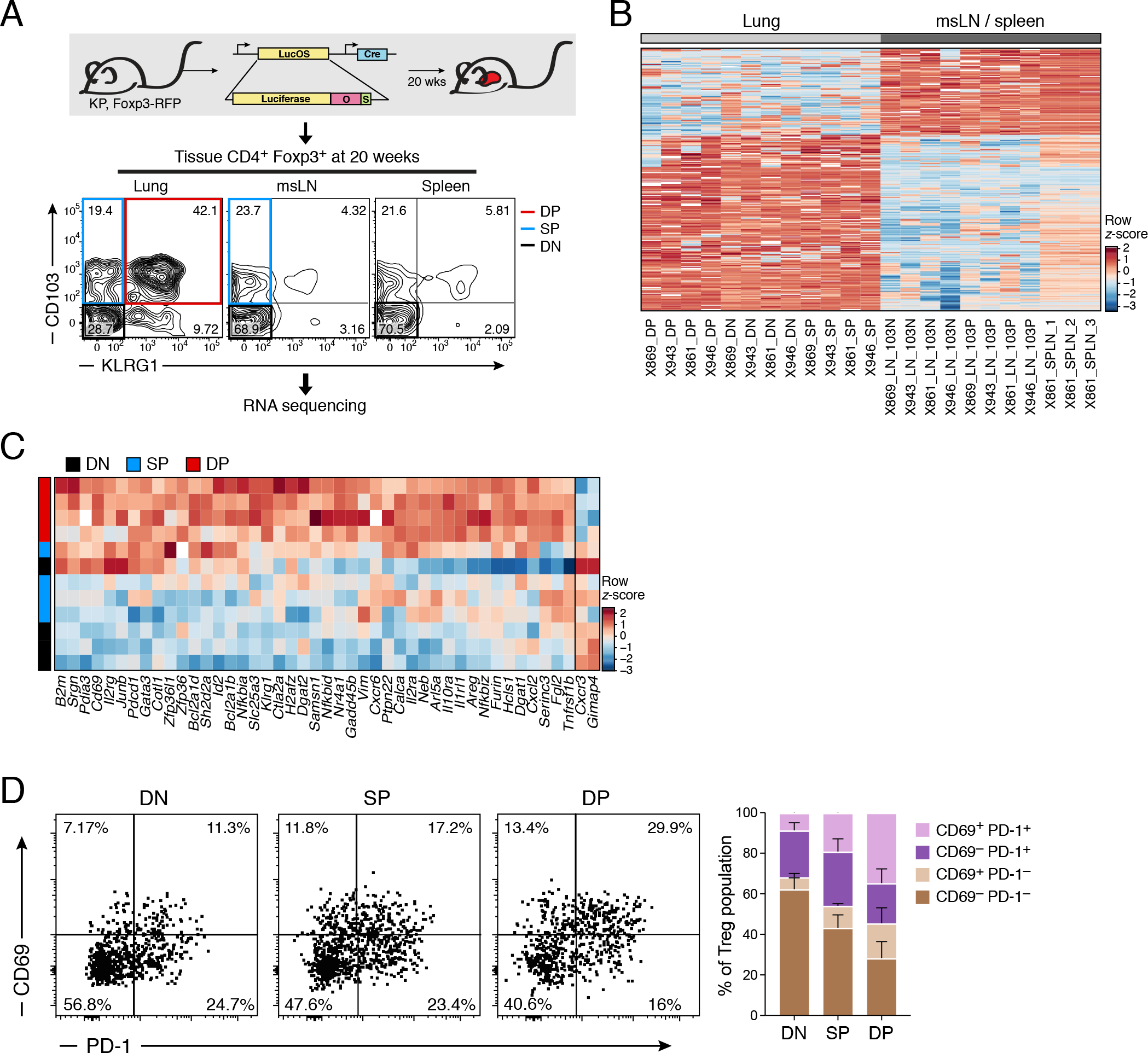
Effector lung T_regs_ from tumor-bearing KP mice bear similarities to activated tissue T_regs_ while demonstrating considerable heterogeneity. **A.** Experiment overview. Top: KP, *Foxp3*^*RFP*^ mice were sacrificed at 20 weeks p.i. Bottom: RNAseq was performed on CD103^−^KLRG1^−^(DN, black), CD103^+^KLRG1^−^(SP, blue), and CD103^+^KLRG1^+^ (DP, red) T_reg_ cells isolated from tumor-bearing lungs, SP and DN T_reg_ cells from the draining mediastinal lymph node (msLN), and DN T_reg_ from one spleen as control. **B.** Gene expression differences (KPLungTR signature genes, |z-score| > 3, |fold change| > 2, **Methods**) between lung (left, gray) vs. msLN/ spleen (right, black) T_regs_ (columns). **C.** 45 gene signature (43 up-regulated, 2 down-regulated) distinguishing DP lung T_regs_ (red) from other populations (black and blue) (**Methods**). **D.**Heterogeneity of CD69 and PD-1 expression among T_reg_ subsets. Representative flow cytometry plots (left) and average cell proportions (right, 3 experiments, each with n=5-6 mice) of CD69 and PD-1 expression among DN, SP, and DP T_regs_. Populations shown are i.v.^neg^CD8^−^CD4^+^Foxp3^+^. Error bars: SEM.

To identify such a program, we bred KP mice to *Foxp3* reporter mice to facilitate isolation and manipulation of T_regs_ from tumor-bearing mice. Using a previously-described method (Anderson et al., 2012), mice were injected with antibody prior to sacrifice to label intravascular cells and distinguish tissue-infiltrating populations. We profiled DP, SP, and DN T_regs_ isolated from the lungs of tumor-bearing KP-*Foxp3*^*RFP*^ mice at 20 weeks post infection (p.i.) with Lenti-LucOS by bulk RNA-Seq (**Figure 1A**, **Methods**). We also profiled SP and DN T_regs_ from matching mediastinal lymph nodes (msLNs) and DN T_regs_ from the spleen of one tumor-bearing mouse for comparison.

The most significant distinction in the data by Independent Component Analysis (ICA) was between lung-infiltrating and peripheral T_regs_ (**Figure S1A**). A 284 gene signature strongly distinguished lung-infiltrating T_regs_ (“KPLungTR signature genes”, **Methods**, **Figure 1B**, **Table S1**), which we confirmed by quantitative RT-PCR (qPCR) of *Pparg1*, *Nr4a1*, *Areg*, and *Gata1*expression (**Figure S1B**). This KPLungTR signature was enriched for signatures of other tissue T_regs_, including T_regs_ in visceral adipose tissue (VAT), colonic lamina propria, and wounded muscle (**Figure S1C**, **Table S2**). Genes upregulated in the KPLungTR signature also included activation, differentiation, and growth factor signaling genes (**Figure S1D**), consistent with prior reports that T_regs_ promote tissue repair (Arpaia et al., 2015; Burzyn et al., 2013). Notably, the signature was enriched for orthologs of genes induced in human colorectal cancer (CRC) and NSCLC-associated T_regs_ (De Simone et al., 2016) (**Figure S1E**), suggesting that lung T_regs_ in human cancer and the KP model have a common “tissue T_reg_” phenotype.

Several lines of evidence further suggest that the DP population is activated. First, genes upregulated and downregulated transiently in activated T_regs_ were differentially expressed in DP *vs*. DN T_regs_ (**Figure S1F**), which may reflect antigen exposure of this T_reg_ population in the tumor microenvironment (van der Veeken et al., 2016). Second, genes upregulated in DP T_regs_ *vs*. all other T_regs_ in tumor-bearing lungs (**Methods**, **Figure 1C**) were associated with T cell activation and putative T_reg_ effector functions (*e.g.*, *Nr4a1*, *Cd69*, *Il1rl1*, *Areg*, *Srgn*, and *Fgl2*). Notably, *Cxcr3*, which has been associated with a T-bet^+^ T_reg_ phenotype specialized to counter Th1 inflammation (Koch et al., 2009; Levine et al., 2017), was downregulated in DP T_regs_ *vs*. SP and DN T_regs_ (**Figure 1C**). The DP T_reg_ phenotype may thus represent an effector cell state different from Cxcr3^+^T-bet^+^ T_regs_.

While the DP subset of lung T_regs_ may be particularly active and an attractive target for immunotherapy, PD-1 and CD69 expression across DN, SP, and DP T_regs_ revealed considerable heterogeneity *within* each subset (**Figure 1D**). In particular, 52% of DP T_regs_ expressed PD-1 and 68% expressed CD69. We thus turned to more fully characterize the variation within T_regs_ in the tumor microenvironment.

### scRNA-Seq reveals heterogeneity within tumor-associated CD4+ T_conv_ cells

We sought to characterize patterns of heterogeneity in tumor-associated CD4^+^ T cells over time to contextualize the diversity of T_reg_ responses in relation to their Foxp3-CD4+ T cell (T_conv_) counterparts. By scRNA-seq we profiled 1,254 T_conv_ and 1,679 T_regs_ sorted from the lungs and msLN of non-tumor bearing KP-*Foxp3*^*GFP*^ mice and tumor-bearing mice at weeks 5, 8, 12, and 20 after tumor induction with Lenti-LucOS (**Figure 2A**,~4 mice per timepoint).

**Figure 2.**
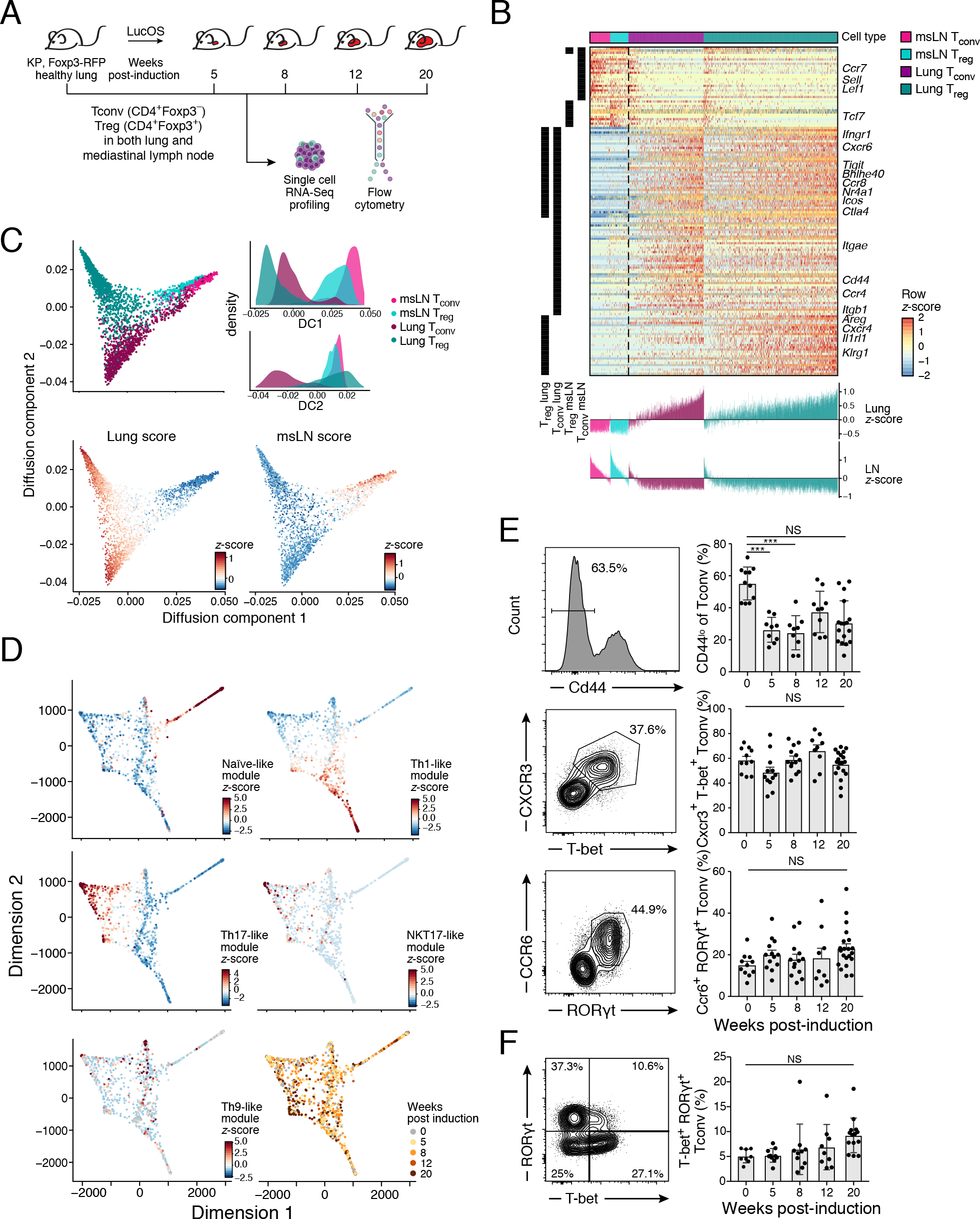
Single-cell RNAseq reveals a distinctive lung CD4^+^ T cell signature and T_conv_ diversity that is stable throughout KP tumor development. **A.**Overview of longitudinal experiment. KP, *Foxp3*^*GFP*^ mice were harvested at the indicated weeks after tumor induction with Lenti-LucOS. 1,254 T_conv_ (i.v.^neg^,Thy1.2^+^CD4^+^Foxp3^−^) and 1,679 T_regs_ (i.v.^neg^,Thy1.2^+^CD4^+^Foxp3^+^) cells from lung and msLN were single-cell sorted and profiled by plate-based scRNAseq. **B.** Shared and lung tissue-specific gene expression program includes genes shared by T_conv_ and T_regs_, and genes unique to each. Genes (rows, row-normalized) differentially-expressed (**Methods**) between cells (columns) from lung (purple, teal) vs. msLN (pink, light blue) for both T_reg_ and T_conv_. Left black bars indicate whether a gene is significantly differentially expressed for T_reg_ and/or T_conv_. Bottom: Each cell’s score (y-axis) for its expression of the corresponding lung and LN signatures, which are different for T_reg_ and T_conv_. Color indicates whether a cell was sorted as a T_reg_ or T_conv_, and tissue of origin. **C.** Lung and msLN cells span a phenotypic continuum, with lung cells showing particular diversity. Diffusion component embedding of all cells (dots), colored by sorted identity and tissue of origin (top left), or by *z*-score of the lung (bottom left) or msLN (bottom right) signatures as in **B**. Top right: distribution of diffusion component (DC) scores for cells from each of the four sorted populations, showing greater range of scores for lung cells. **D-F.** Lung T_conv_ subsets expressed programs associated with naive/ central memory T, Th17, Th1, Th9, and NKT17 cells. **D.** Two-dimensional force-directed layout embedding of the first four diffusion components of all lung resident T_conv_ cells (**Methods**), with cells colored by expression z-score for the indicated gene module, or by timepoint after tumor induction (bottom right). **E-F.** Left: Representative flow cytometry plots demonstrating naive/ central memory (E, top), Th1 (E, middle), Th17 (E, bottom), and Th1/Th17 (F, T-bet^+^RORγt^+^) CD4^+^ T cell populations. Right: Corresponding barplots showing the percentage (y-axis) of the indicated T_conv_ (i.v.^neg^CD8^−^CD4^+^Foxp3^−^) subset throughout tumor development (x-axis) across 2-3 experiments (dot: one mouse). Error bars: SEM. ***p<0.001, Tukey’s multiple comparisons test. NS: non-significant.

The tissue-specific expression program partitioned into genes shared by lung infiltrating T_conv_ and T_regs_, and genes uniquely upregulated in each (**Figure 2B**, **Table S3**). For example, lung-infiltrating T_regs_ expressed high levels of *Il1rl1, Cxcr4, Areg*, and *Klrg1*, while T_conv_ cells expressed *Cd44, Ccr4* and *Itgb1* (**Figure 2B**). Genes from the KPLungTR signature and from a recently described trajectory of tissue-resident T_regs_ (Miragaia et al. 2017) were both differentially expressed in the scRNA-seq profiles (**Figure S2A**).

Both the lung and msLN cells spanned a phenotypic continuum, with the lung cells showing particular diversity (**Figure 2C, S2B**, DC1 p < 10^−13^; DC2 p < 10^−16^, Levene’s test). The spectrum of cell states was apparent when scoring for the expression of lung T_conv_ or T_reg_ signatures, and when cells were arranged along diffusion components that describe their tissue-specific expression program (**Figure 2C**). Both T_regs_ and T_conv_ in the msLN expressed genes associated with a naive or central memory phenotype, including *Lef1*, *Sell*, and *Ccr7* (**Figure 2B, S2C**). Conversely, cells were more activated in the lung (**Figure 2B**). Subsets of lung T_conv_ and T_reg_ cells that scored highly for the msLN signature also expressed genes associated with TCR signaling, including *Nr4a1* and *Junb*, suggesting that they may be recently activated (**Figure 2C**, **S2C**). Lung-infiltrating T_conv_ and T_reg_ cells that scored highly for the respective lung signature may represent cells that were more tissue-adapted or localized to a particular region of the lung.

### Lung T_conv_ subsets remain in stable proportions throughout tumor development

Lung T_conv_ subsets expressed programs associated with different CD4^+^ T cell subsets, including naïve T, Th17, Th1, Th9 and NKT17 cells (**Methods**, **Figure 2D-E**), whose proportions remained largely stable over time. Within Th1 cells, a subset expressed *Eomes* and *Gzmk*, which may reflect cytolytic function, and *Cxcr3* and *Ccr5*, which promote antigen-specific CD4^+^ T cell recruitment to lungs during respiratory virus infection (Kohlmeier et al., 2009) (**Figure S2D**). Some of the Th17-like cells expressed *Zbtb16*, a marker for NKT cells, and also scored highly for a gene module that includes genes associated with natural killer T17 (NKT17) cells, such as *Blk* and *Gpr114* (**Figure S2E**) (Engel et al., 2016). Furthermore, these cells had lower expression of CD4 than other T_conv_ (**Figure S2F**) and did not express TCR chains associated with γδ T cells (**Table S5**). We found little evidence of Th2-like cells, despite their role in lung inflammation in other settings (Walker and McKenzie, 2018), but did observe a small population of Th9-like cells expressing *Il9r*, *Il4*, and *Il1rl1*, which have been implicated in driving anti-tumor immune responses (Végran et al., 2015) (**Figure S2G**). Finally, we identified a population that scored highly for both the Th1 and the Th17 modules. We validated the presence of cells expressing both RORγt and T-bet (**Figure 2F**); such cells have been described as a plastic, Th17-derived population in other pathogenic states (Lee et al., 2012, 2009; Wang et al., 2014). The overall expression of the gene modules associated with these T_conv_ subsets showed subtle variation over time by scRNA-Seq (**Figure S2H-I**), but the relative cell proportions measured by flow cytometry remained stable during tumor development (**Figure 2E-F**).

### A RORγt^+^ T_reg_ population is present throughout tumor development and may have shared clonal origin with Th17 T_conv_ cells

Lung-infiltrating T_regs_ expressed several gene modules with similar features to those in transcriptional signatures of previously-described T_reg_ subsets (**Figure S3A**). For example, Module 18 includes genes that characterize a resting, or central, T_reg_ (rT_reg_) phenotype, such as *Sell*, *Ccr7*, and *Tcf7* (Campbell, 2015; Li and Rudensky, 2016), whereas Module 13 identified a T_reg_ population expressing *Rorc* and *Il17a* (**Figure 3A**, **S3A**), reminiscent of Th17-like T_regs_ (Tr17), a subset with immunosuppressive activity directed at Th17 responses (Kim et al., 2017). We validated this population by flow cytometry and found that RORγt^+^ T_regs_ comprise roughly 10% of lung-infiltrating T_regs_ throughout tumor progression (**Figure 3B**). The Tr17-like cells represented a distinct state among lung T_regs_ and the expression of Tr17-associated genes was inversely correlated with the expression of genes previously identified in lung-resident T_regs_, including KLRG1 (**Figure 3C-D**). Additionally, whereas *Ccr6* expression within the T_conv_ was restricted to Th17 cells (**Figure 2E**), *Ccr6* was expressed in multiple T_reg_ subsets (**Figure S3B**), consistent with previous findings (Yamazaki et al., 2008), which may result in the localization of different T_reg_ subsets to common sites in the lung.

**Figure 3.**
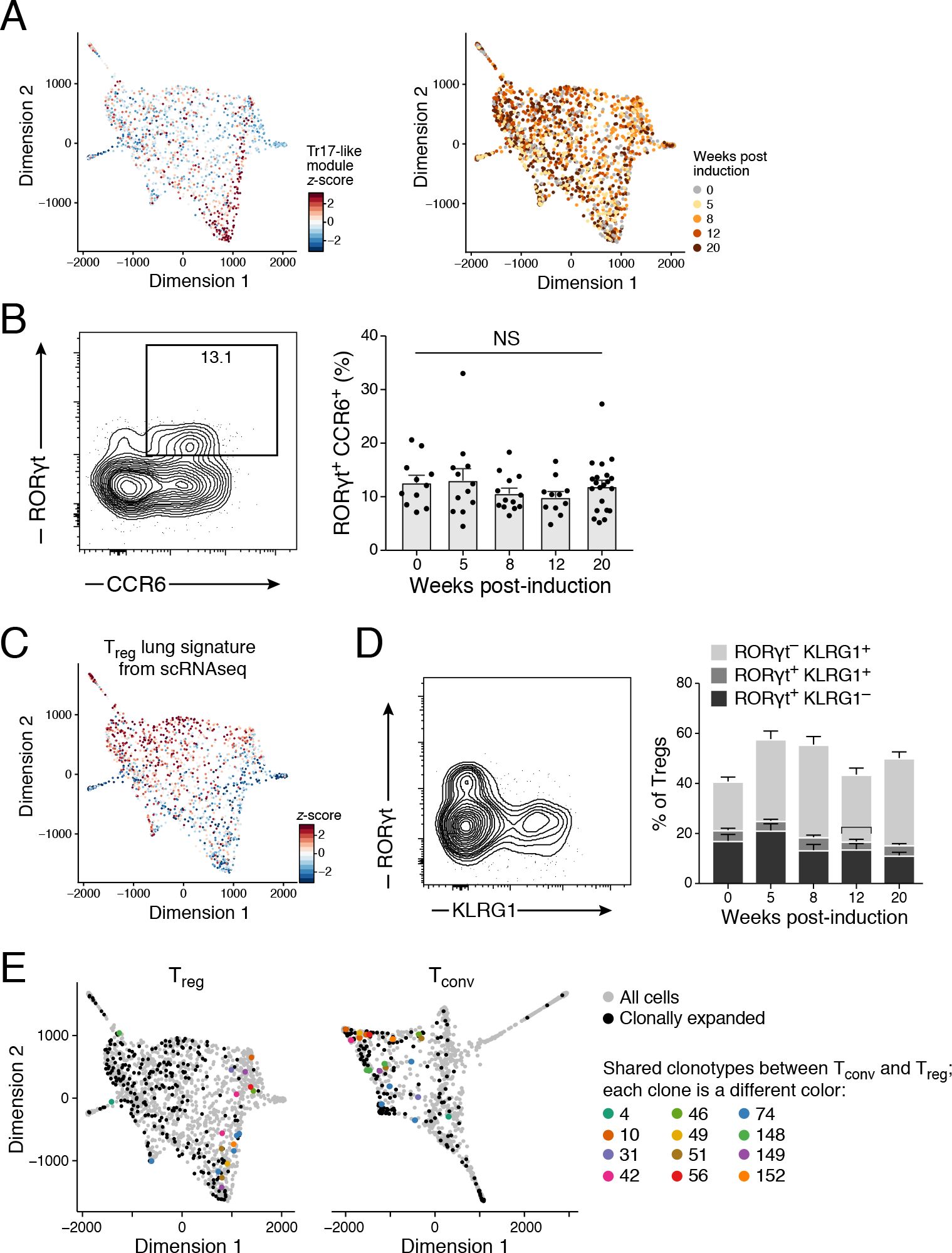
A Th17-like T_reg_ population is present throughout tumor development and may have shared clonal origin with T_conv_ cells. **A-B.**Cells expressing a Tr17-like program are present throughout tumor development. **A.** Two-dimensional force-directed layout embedding of the first six diffusion components of all lung-derived T_regs_ where each cell (dot) is colored by the normalized average gene expression (z-score) of the genes in module 13, which represents *Rorc*^+^ T_regs_ (left) or by timepoint after induction (right). **B.**Left: Representative flow cytometry plot demonstrating RORγt^+^CCR6^+^ T_regs_(i.v.^neg^CD8^−^CD4^+^Foxp3^+^). Right: Percentage of T_regs_ that are RORγt^+^CCR6^+^ (y-axis) across tumor development (x-axis) across 2-3 experiments. Error bars: SEM. NS: non-significant, Tukey’s multiple comparisons test. **C-D.**Tr17-like and T_reg_ programs are inversely correlated. **C.** Two-dimensional force-directed layout embedding of all lung-derived T_regs_ as in A, with each cell (dot) colored by normalized average gene expression (*z*-score) of the genes upregulated in lung vs. msLN T_regs_ (as in **Figure 2B)**. **D.** Left: Representative flow cytometry plot of T_reg_ (i.v.^neg^CD8^−^CD4^+^Foxp3^+^) expression of RORγt and KLRG1 (left). Right: Percentage of T_regs_ that are RORγt^+^KLRG1^+^, RORγt^+^KLRG1^−^, and RORγt^−^KLRG1^+^ across tumor development (x-axis) across 2-3 experiments (dot = one mouse). Error bars: SEM. **E.**Shared clonotypes between T_reg_ and T_conv_ are predominantly in Tr-17 like and Th17-like cells. Two-dimensional force-directed layout embedding of lung-resident T_regs_ (left, as in **A**) and T_conv_ (right, as in **Figure 2D**) with each cell colored by clonal analysis. Grey: not clonal at our resolution or no TCR was reconstructed. Black: cells that share a TCR with at least one other cell. Color: Shared clones between T_reg_ and T_conv_, with numeric identifiers.

Remarkably, shared clonotypes between T_reg_ and T_conv_ cells were predominantly Tr17-like and Th17-like cells, respectively. Specifically, based on paired-chain T cell receptor (TCR) sequences of profiled cells (**Figure S3B**, **Methods**, **Table S5**), 12 TCR clonotypes were shared across T_reg_ and T_conv_ cells. Indeed, dedicated TCR profiling of T_regs_ and T_conv_ from KP mice with advanced disease showed that ~5% of T_reg_ clones were shared with T_conv_ on average in advanced disease (**Figure S3C**). Of the 19 T_regs_ and 20 T_conv_ cells belonging to the 12 TCR clonotypes shared between T_conv_ and T_reg_, the T_reg_ cells were predominantly of the Tr17-like phenotype (13 of 19 T_regs_ had a *z*-score > 1.5 in the Tr17-like Module, hypergeometric p-value < 10^−5^, **Figure 3F**, **S3D**). The T_conv_ cells were also predominantly of the Th17 phenotype, although this was not a significant enrichment . 67 out of 178 identified T_conv_ clones were of the Th17 phenotype (hypergeometric p = 0.68), of which 8 were clonotypes shared with T_regs_, (**Figure 3F**). Thus, Tr17 differentiation may reflect a shared clonal origin with Th17 cells.

### An effector-like T_reg_ phenotype becomes predominant during tumor development

In contrast to Tr17-like cells, where a program was expressed by a fixed proportion of cells during tumor development, other T_reg_ programs changed in prominence throughout tumor development (**Figure 4A**). For example, there was decreased expression of Modules 1, 3, 8, and 9, which mark cycling cells, after 8 weeks (**Figure 4A**), corresponding to a decline in Ki67 expression on T_regs_ (**Figure 4B**). Two other programs also changed over time, reflecting an interferon response and a T effector program (**Figure 4A**).

**Figure 4.**
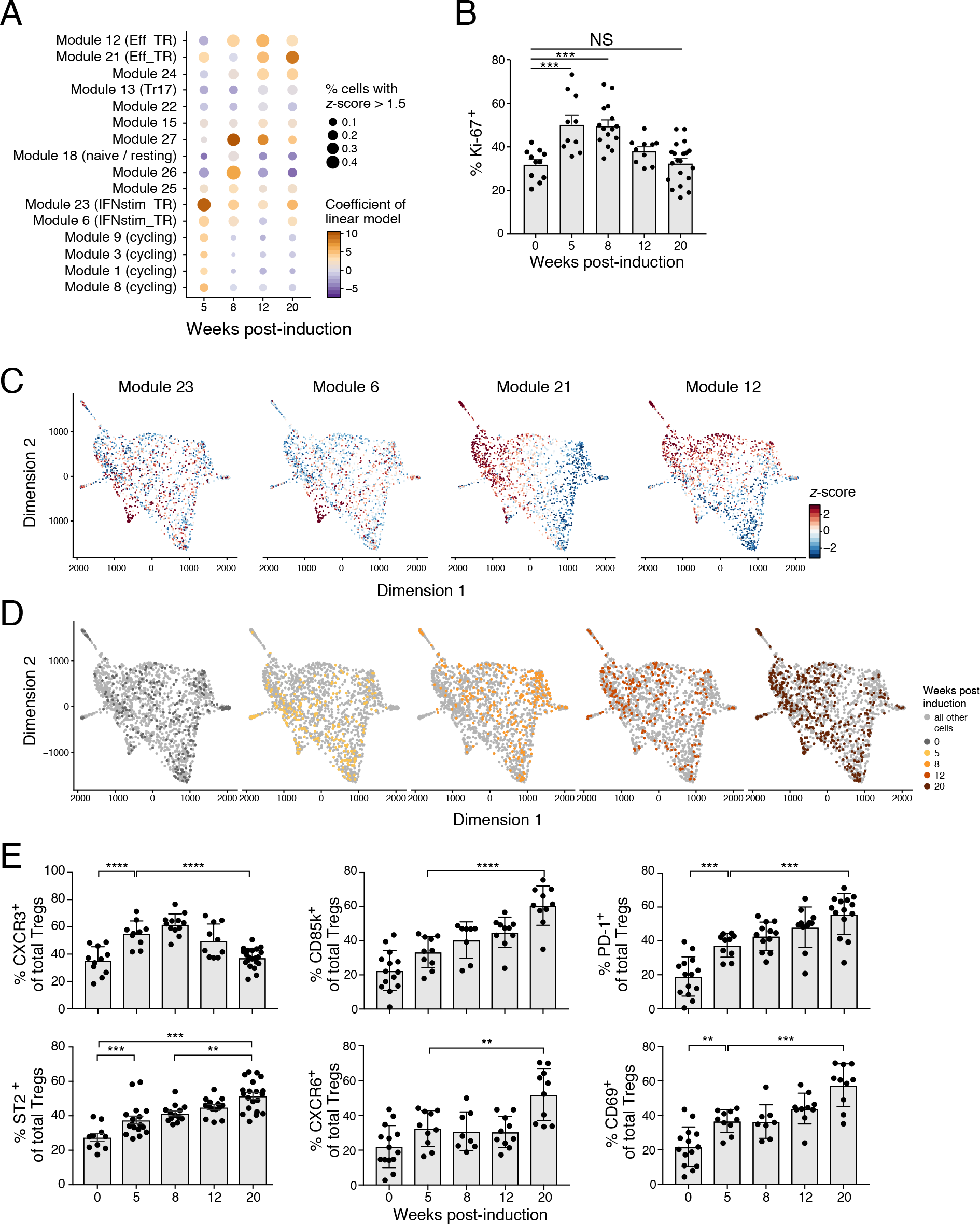
An effector T_reg_ phenotype becomes dominant during tumor development. **A.** Changes in prominence of cycling, interferon-stimulated, and T_reg_ effector programs with tumor development. Linear regression analysis of module expression z-scores as a function of time since tumor initiation, where non-tumor bearing lung is the reference for the timepoint covariate. Dot plot shows for each module (row) and timepoint (column) the coefficients of the timepoint covariate of the regression (color), and the percentage of cells with a *z*-score > 1.5 (dot size). Brown/ blue: increased/ decreased expression over time compared to non-tumor bearing lung. **B.** T_reg_ proliferation peaks early in tumor development. The percentage of Ki-67^+^ T_regs_(y-axis) throughout KP tumor development (x-axis) from 2-3 experiments (dot = one mouse). Error bars: SEM. ***p <0.001, Tukey’s multiple comparisons test. NS: non-significant. **C-E.**An interferon and an effector program peak early and late in tumor development, respectively. **C-D.** Two-dimensional force-directed layout embedding of all lung-infiltrating T_regs_ (as in **Figure 3A**) colored by normalized signature z-score for the IFNstim_TR modules (C, Modules 6 and 23) and the Eff_TR modules (C, Modules 12 and 21), or timepoint after tumor induction (D). **E.** Percentage of T_regs_ expressing the indicated protein (y-axis) throughout KP tumor development (x-axis) from 2-3 experiments (dot: one mouse). Error bars: SEM. **p<0.01, ***p<0.001, ****p<0.0001, Tukey’s multiple comparisons test.

The interferon program (“IFNstim_TR”) was characterized by the expression of Modules 6 and 23 (**Figure 4C**), which included many interferon-stimulated genes (ISGs) downstream of either type I or II interferon (IFN) signaling, including *Stat1*, guanylate binding protein genes (GBPs), type I interferon-specific genes (*e.g.*, oligoadenylate synthetase family members), and IFNγ-specific genes (*e.g*., *Irf1, Irf9*) (Der et al., 1998). 28 genes from the IFNstim_TR program were significantly downregulated by T_regs_ during tumor progression (**Figure S4B**). IFNγ promotes a Tbet^+^CXCR3^+^ Th1-like T_reg_ cell population that can suppress Th1 responses (Hall et al., 2012; Koch et al., 2009, 2012). Neither *Cxcr3* nor *Tbx21* are IFNstim_TR genes, but IFNstim_TR expression was correlated with *Tbx21* expression (**Figure S4C**). Moreover, the program was enriched for genes expressed by lymphoid tissue T_regs_ and genes downregulated in DP T_regs_ (**Figure S4D**), which include *Cxcr3*. IFNstim_TR expression may thus reflect recent arrival to the lung, consistent with its presence early in tumor development.

The T effector program (“Eff_TR”) was characterized by the expression of Modules 12 and 21 (**Figure 4C**), which were enriched for genes in the DP signature (p-value ≤ 10^−25^, **Figure S4E**) and genes upregulated in T_regs_ from mouse non-lymphoid tissues and human breast cancer, NSCLC, and CRC (De Simone et al., 2016; Guo et al., 2018; Magnuson et al., 2018; Miragaia et al., 2017; Plitas et al., 2016; Zheng et al., 2017) (**Figure S4D**), confirming the distinct expression profile we had previously identified in the DP T_reg_ subpopulation.

The interferon and effector programs represented independent phenotypes of T_regs_ within each timepoint but followed opposite patterns over time: expression of IFNstim_TR genes was highest in cells from week 5 and declined thereafter, while expression of Eff_TR genes increased and remained elevated (**Figure 4A, D**). This temporal transition was also highlighted when testing for individual temporally varying genes: *Cxcr3* expression decreased with time, and *Pdcd1* and *Lilrb4* (Module 21) increased in expression during tumor development (**Figure S4F**), consistent with down-regulation of *Cxcr3* in DP T_reg_ cells (**Figure 1D**). More generally, Eff_TR genes were upregulated in DP T_regs_ compared to DN T_regs_ in mice with late-stage tumor burden, whereas IFNstim_TR genes were significantly downregulated (**Figure S4G**). We confirmed that protein levels of Cxcr3 decreased, and proteins encoded by Eff_TR genes, including CD85k, CD69, CXCR6, PD-1 and ST2, increased during tumor progression (**Figure 4E**).

Taken together, our data suggest that tumor progression may be associated with a shift from a T_reg_ cell phenotype specialized for responding to Th1 inflammation to an effector T_reg_ cell population. In particular, we hypothesized that the strong immunosuppression associated with the late-stage tumor environment may be a result of the emergence and stabilization of cells with the Eff_TR phenotype.

### ST2 is upregulated on effector T_regs_ in mice bearing advanced lung tumors

We reasoned that *Il1rl1*, an Eff_TR gene that encodes the interleukin 33 (IL-33) receptor ST2, may highlight a pathway that could be targeted to alter longitudinal changes in T_reg_ cell phenotype and prevent the accumulation of effector T_regs_ in advanced tumors. First, *Il1rl1*/ ST2 levels tracked with the effector T_reg_ phenotype; *Il1rl1* is a member of Module 21 in the Eff_TR program and ST2 was most highly expressed in DP lung T_regs_ (**Figure 5A**), and its expression in T_regs_ increased during tumor development (**Figure 4D**). Moreover, ST2 was expressed by ~40% of lung T_regs_ *vs*. ~10% of T_regs_ in the msLN, and <5% of T_conv_ cells in the lung in late-stage tumor-bearing mice (**Figure 5B**). Second, T_reg_ cells from tumor-bearing KP, LucOS-infected mice expressed both the membrane-bound and soluble isoforms of ST2 (**Figure 5C**); soluble ST2 (sST2) is thought to diminish ST2 signaling through sequestration of IL-33, the only known ligand of ST2 and an alarmin that recruits immune cells to sites of tissue damage (Cayrol and Girard, 2014). Finally, IL-33 was highly expressed in normal lung, and in early and late lung adenocarcinomas in the KP model (**Figure 5D**). In normal lung, IL-33 was predominantly expressed on surfactant protein C (SPC)-expressing type II epithelial cells (**Figure S5**). We thus hypothesized that ST2 may be a critical mediator of T_reg_ cell function in the lung tumor environment.

**Figure 5.**
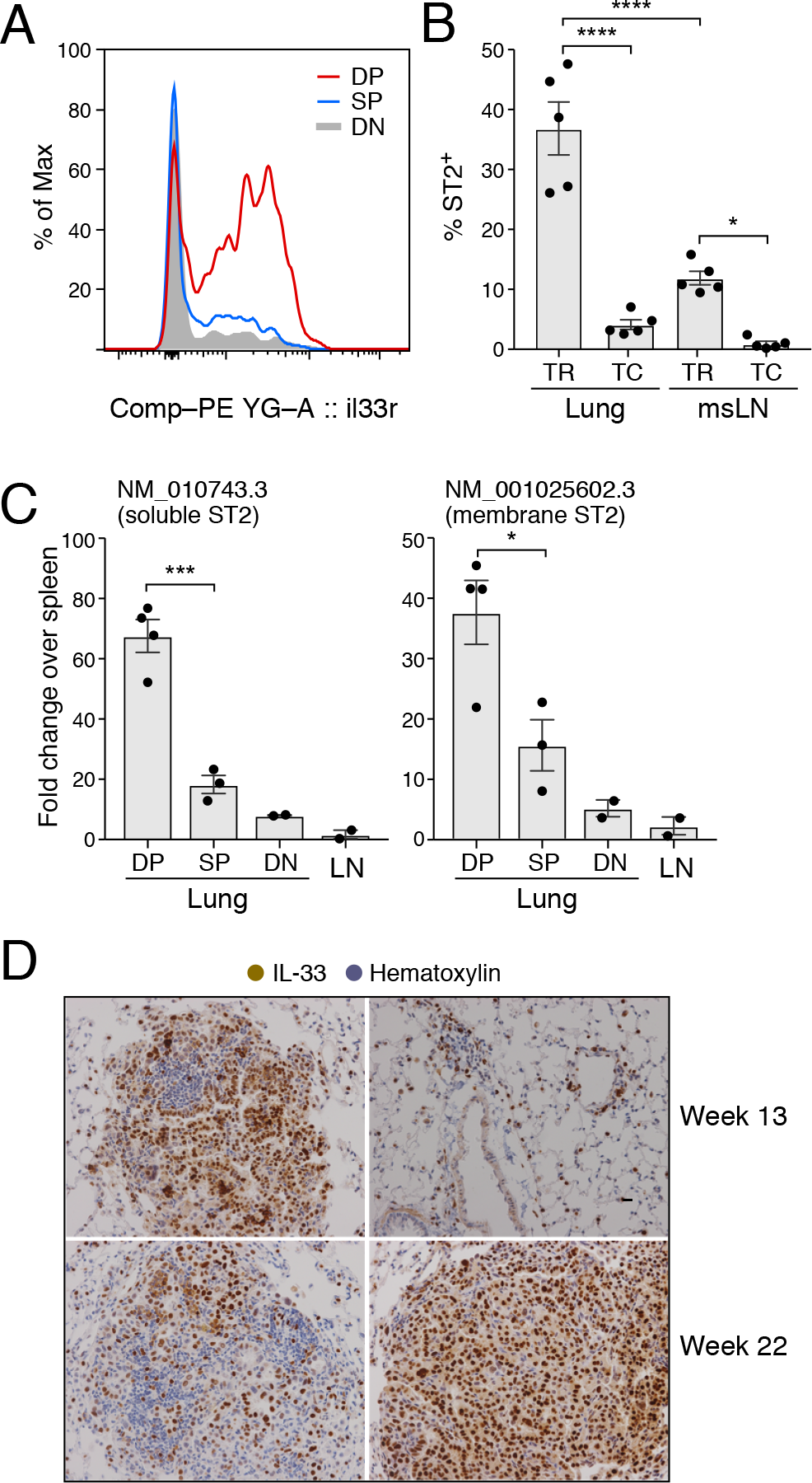
ST2 is upregulated in terminally-differentiated T_regs_ in lung tumor-bearing mice. **A.** ST2 is most highly expressed in DP lung T_regs_. Representative distributions of ST2 expression on CD103^−^KLRG1^−^(DN, grey), CD103^+^KLRG1^−^(SP, blue), and CD103^+^KLRG1^+^ (DP, red) T_regs_ isolated from tumor-bearing lungs. **B.**Lung T_regs_ are enriched for ST2^+^ cells in late-stage tumors. Percent ST2^+^ (y-axis) among lung and msLN T_regs_ (i.v.^neg^CD4^+^Foxp3^+^) and T_conv_ cells (i.v.^neg^CD4^+^Foxp3^−^) (x-axis) from tumor-bearing LucOS mice at week 20 p.i. as measured by flow cytometry. ****p <0.0001, *p <0.05, Tukey’s multiple comparisons test. **C.**T_regs_ from tumor-bearing mice express both the membrane-bound and soluble isoforms of ST2. Relative expression (y-axis, 2^−ΔΔCt^, qRT-PCR, with splenic T_reg_ expression as control) of NM_001025602.3 (left, *Il1rl1* transcript variant 1 encoding membrane-bound ST2) and NM_010743.3 (right, *Il1rl1* transcript variant 2 encoding soluble ST2) in DP, SP, and DN lung T_regs_ and SP and DN msLN T_regs_ (x-axis) (dot: one mouse). Error bars: SEM. ***p <0.001, *p <0.05, Tukey’s multiple comparisons test. **D.** IL-33 is highly expressed in lung adenocarcinoma. Immunohistochemical staining of tumor-bearing lungs from KP mice at weeks 13 and 22 p.i. with Lenti-LucOS. Two representative images are shown per timepoint.

### Recombinant IL-33 treatment increases effector T_regs_ in tumor-bearing lungs

To determine the effect of IL-33 on the immune microenvironment of tumors, we administered recombinant mouse IL-33 (rIL-33) intratracheally to tumor-bearing KP, Lenti-LucOS-infected mice (**Figure 6A**). Consistent with prior reports (Kondo et al., 2008; Schmitz et al., 2005), rIL-33 induced significant inflammatory infiltration and epithelial thickening in tumors and throughout the lung (**Figure 6B**). rIL-33-treated mice had greater numbers of eosinophils (**Figure 6C**) and CD4^+^ and CD8^+^ T cells per lung weight (**Figure 6D**), although the proportion of tumor-specific, SIINFEKL tetramer-positive cells among CD8+ T cells was unchanged (**Figure 6E**). We observed similar inflammation in non-tumor bearing wild-type mice treated with rIL-33 (data not shown). CD4^+^ T cells in rIL-33-treated mice had an increased proportion of T_regs_ (**Figure 6F**), of which 64% were DP compared to 34% in PBS-treated controls, with proportionally fewer SP and DN T_regs_ (**Figure 6G**). rIL-33 treatment of ST2-deficient mice failed to elicit the same change in the proportion of T_regs_, which was similar to that of untreated, wild-type mice (**Figure S6**). Taken together, rIL-33 administration is sufficient to drive both a major increase in the lung T_reg_ population in general, and to promote an increase in effector T_regs_ cells in particular.

**Figure 6.**
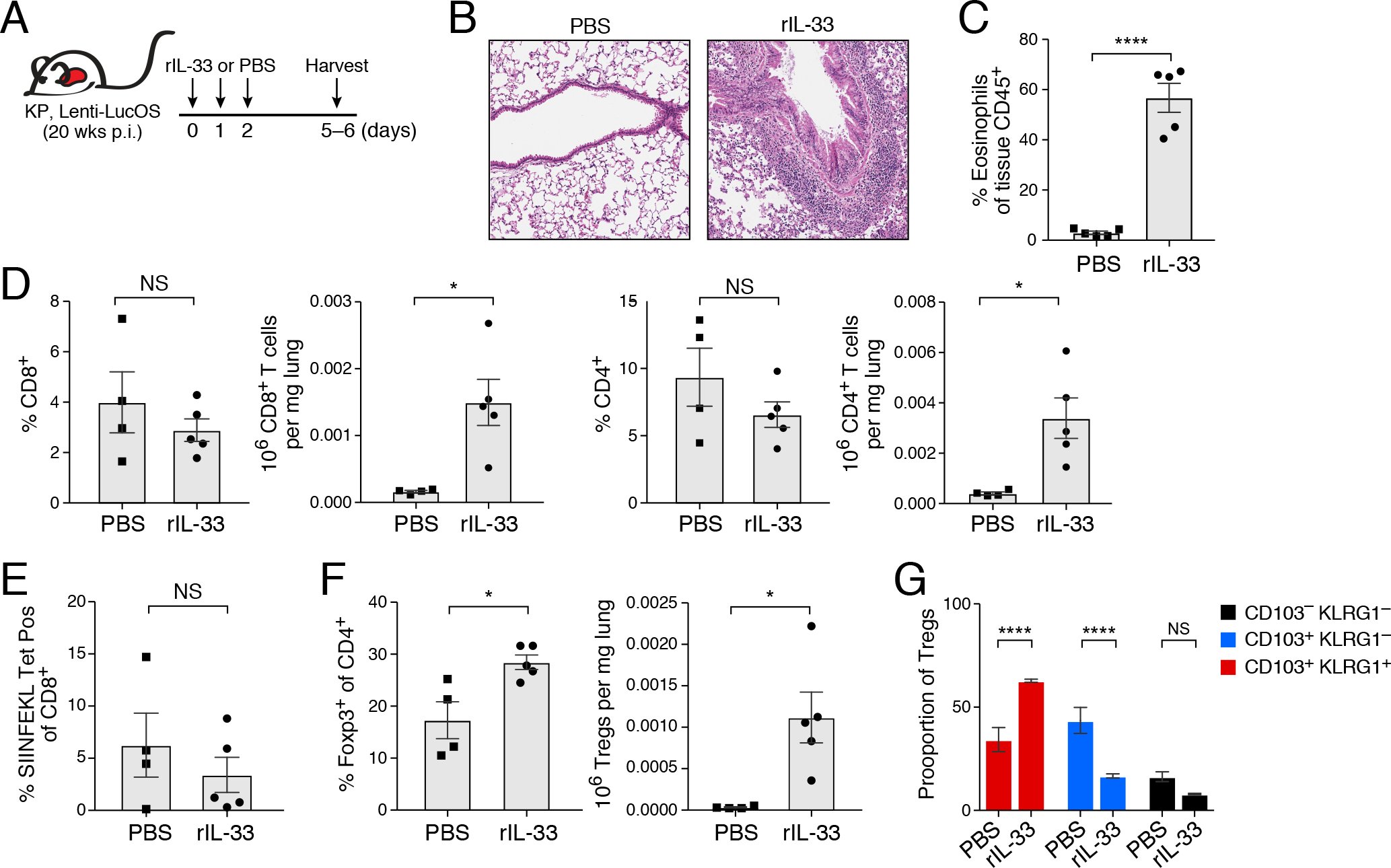
rIL-33 is sufficient to promote an increase in effector T_regs_ in tumor-bearing lungs. **A.** Experimental overview. Recombinant IL-33 (rIL-33) or PBS control were administered to late-stage, tumor-bearing KP mice. All rIL-33 experiments are representative of 2-3 separate experiments, each with n=4-5 mice per group. **B-D.** rIL-33 induced inflammatory infiltration and epithelial thickening. **B.** Representative hematoxylin and eosin (H&E)-stained histological images of control (left) and rIL-33-treated (right) lungs at 10X magnification. **C.** Proportion of eosinophils (y-axis, i.v.^neg^CD45.2^+^CD11c^−/low^, SiglecF^+^) of i.v.^neg^CD45^+^ lung cells from control and rIL-33-treated mice. Data is representative of 2 independent experiments. Error bars: SEM. ****p < 0.0001, two-tailed Student’s t test. **D.** Proportions (y-axis, left) and absolute numbers (y-axis, right) of lung CD8^+^ and CD4^+^ T cells of i.v.^neg^ cells in control and rIL-33-treated mice (x-axis). Error bars: SEM. *p = 0.01, two-tailed Student’s t test. **E.** No change in proportion of SIINFEKL tetramer-positive CD8^+^ T cells. Percentage of SIINFEKL/Kb tetramer-positive cells out of lung i.v.^neg^CD8^+^ T cells (y-axis) in control and rIL-33-treated mice (x-axis). Error bars: SEM. NS: non-significant, Tukey’s multiple comparisons test. **F.** Increase in T_reg_ proportions in rIL-33-treated mice. Proportion (y-axis, left) and absolute number (y-axis, right) of T_reg_ cells out of i.v.^neg^CD4^+^ lung T cells in control and rIL-33-treated mice (x-axis). Error bars: SEM. *p = 0.02, two-tailed Student’s t test. **G.**Reduced changes in T_reg_ proportions in rIL-33-treated, ST2-deficient mice. Percent of cells (y-axis) that are CD103^−^KLRG1^−^(DN, black), CD103^+^KLRG1^−^(SP, blue), or CD103^+^KLRG1^+^ (DP, red) out of T_reg_ cells from tumor-bearing lungs of control and rIL-33-treated mice. Error bars: SEM. ****p < 0.0001, Sidak’s multiple comparisons test. NS: non-significant.

### T_reg_-specific ST2 is required for the increase in effector T_regs_ during tumor progression

To test whether ST2 signaling on T_regs_ was necessary for the development of a robust effector T_reg_ cell response in tumors, we studied the effects of T_reg_-specific *Il1rl1* deletion. We used a modified version of the KP model wherein FlpO recombinase drives expression of oncogenic K-ras and loss of p53 (KPfrt: *FSF-Kras^G12D^, p53^frt/frt^*), which allowed us to use the Cre-lox system to study T_reg_-specific *Il1rl1* deletion. We crossed KPfrt mice to *Foxp3*^*YFP-Cre*^ and *Il1rl1*^*fl/fl*^ mice to model lung adenocarcinoma development in the setting of T_reg_-specific ST2 deficiency (**Figure 7A**). We infected the mice with a lentivirus expressing FlpO recombinase and GFP fused to Ova and SIYRGYYL (FlpO-GFP-OS) in order to induce tumors that would express the same strong T cell antigens as those in the Lenti-LucOS model.

**Figure 7.**
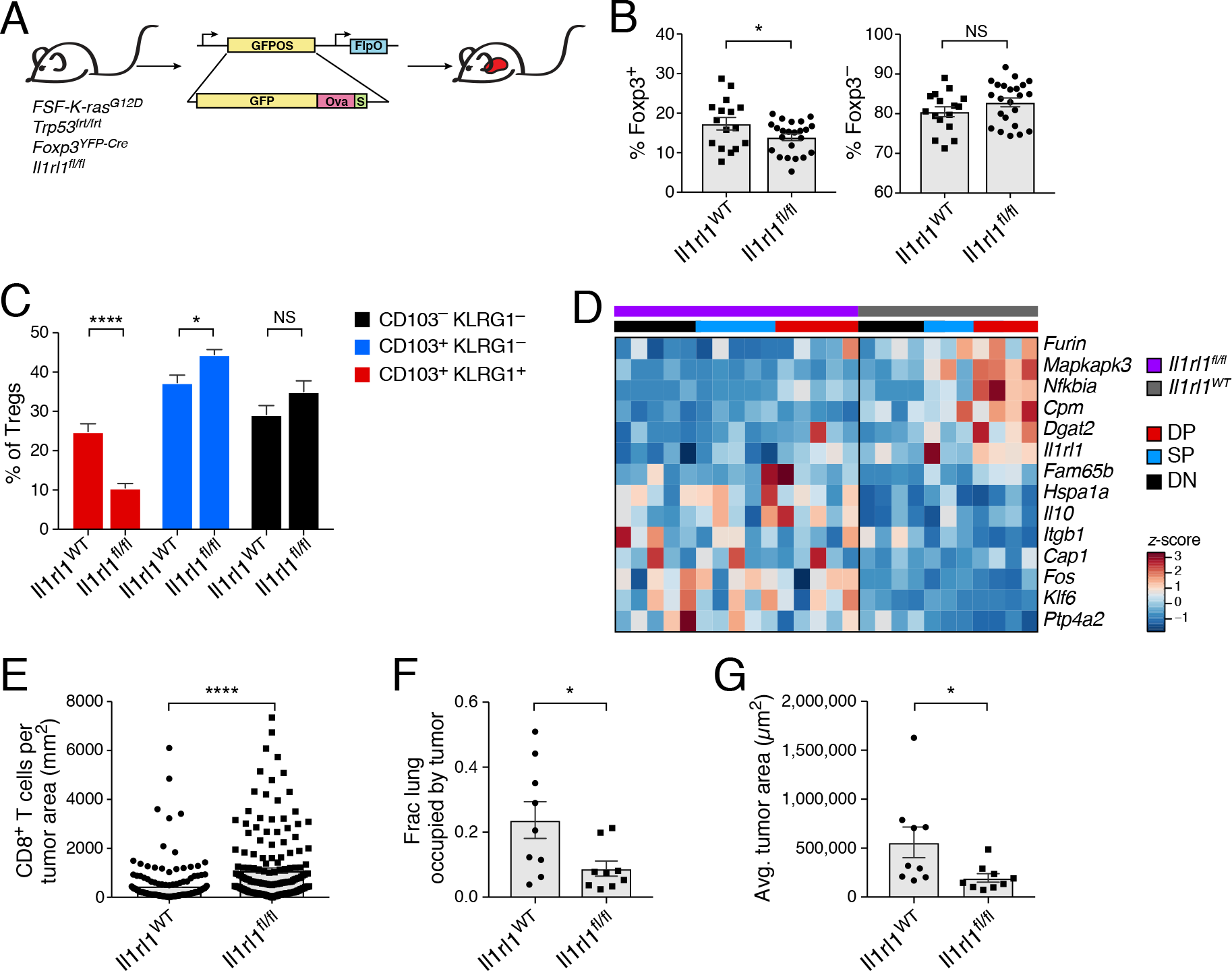
T_reg_-specific ST2 ablation impairs expansion of effector T_reg_ cells and enhances CD8^+^ T cell infiltration of tumors. **A.** Experiment overview. KPfrt, *Foxp3*^*YFP-Cre*^(“*Il1rl1^WT^*”)and KPfrt, *Foxp3*^*YFP-Cre*^, *Il1rl1*^*fl/fl*^ (“*Il1rl1^fl/fl^*”) mice were infected with Lenti-FlpO-GFP-OS. **B-C.** Changes in T_regs_ and their subsets in *Il1rl1*^*fl/fl*^ mice with advanced lung tumors. **B.** Percent of Foxp3^+^ (y-axis, left) and of Foxp3^−^(y-axis, right) of i.v.^neg^CD4^+^ lung cells in KPfrt, *Foxp3*^*YFP-Cre*^ vs. KPfrt, *Foxp3*^*YFP-Cre*^, *Il1rl1*^*fl/fl*^ mice at 24-25 weeks p.i across 3 experiments, each with n=3-5 mice per group. Error bars: SEM. *p<0.05, two-tailed Student’s t test. NS: non-significant. **C.** Percent of CD103^−^KLRG1^−^(DN, black), CD103^+^KLRG1^−^(SP, blue), and CD103^+^KLRG1^+^ (DP, red) out of T_regs_ isolated from the tumor-bearing lungs of KPfrt, *Foxp3*^*YFP-Cre*^ vs. KPfrt, *Foxp3*^*YFP-Cre*^, *Il1rl1*^*fl/fl*^ mice across 3 experiments, each with n=3-5 mice per group. Error bars: SEM. ****p < 0.0001, *p < 0.05, Sidak’s multiple comparisons test. NS: non-significant. **D.** Expression signature distinguishing *Il1rl1*^*WT*^ from ST2-deficient T_regs_ from tumor-bearing mice. Row-normalized expression (z-score) of select signature genes (rows, **Methods**) across CD103^−^KLRG1^−^(DN, black), CD103^+^KLRG1^−^(SP, blue), and CD103^+^KLRG1^+^ (DP, red) T_regs_ (columns, lower color bar) from KPfrt, *Foxp3*^*YFP-Cre*^(gray) vs. KPfrt, *Foxp3*^*YFP-Cre*^, *Il1rl1*^*fl/fl*^ (purple) mice. **E.** Increased CD8+ T cell infiltration in mice with T_reg_-specific ST2 deficiency. Number of CD8^+^ cells per tumor area (y-axis) in pooled tumors from KPfrt, *Foxp3*^*YFP-Cre*^ and KPfrt, *Foxp3*^*YFP-Cre*^, *Il1rl1*^*fl/fl*^ mice across two experiments, with n=4-5 mice per group. CD8 was measured by immunohistochemical (IHC) staining of histological cross-sections of tumor-bearing lungs. Error bars: SEM. ****p<0.0001, Mann-Whitney test. **F-G.** Reduced tumor burden in mice with T_reg_-specific ST2 deficiency. Percent of total lung occupied by tumor (F, y-axis) and average tumor size (G, y-axis, μm^2^) in KPfrt, *Foxp3*^*YFP-Cre*^ vs. KPfrt, *Foxp3*^*YFP-Cre*^, *Il1rl1*^*fl/fl*^ mice in histological cross-sections of tumor-bearing lungs across two experiments, with n=4-5 mice per group. Error bars: SEM. *p=0.0315 (F), 0.0106 (G), Mann-Whitney test.

Early-stage KPfrt, *Foxp3*^*YFP-Cre*^, *Il1rl1*^*fl/fl*^ mice did not differ from KPfrt, *Foxp3*^*YFP-Cre*^ mice in the fraction of CD4^+^ T cells that were T_conv_ or T_reg_ cells, but late in tumor progression there was a slight reduction in the proportion of T_reg_ cells (**Figure 7B** and **S7A**), a significantly lower proportion of DP T_regs_, and a higher proportion of SP cells (**Figure 7C**). Expression profiles of DP, SP, and DN T_regs_ from KPfrt, *Foxp3*^*YFP-Cre*^, *Il1rl1*^*fl/fl*^ and KPfrt, *Foxp3*^*YFP-Cre*^ control mice identified an expression signature lower in ST2-deficient vs. wild-type T_regs_, where it was highest among wild-type DP T_regs_ (**Figure 7D, S7D**). The signature was enriched for KPLungTR and DP signature genes, including *Dgat2, Furin* and *Nfkbia*, as well as for genes upregulated by T_regs_ in human NSCLC (**Figure 7E**, **S7B, C**). ST2-deficient T_regs_ also showed higher expression of some genes, including *Itgb1*, *Il10*, *Klf6*, and *Fos* (**Figure 7E**), suggesting that they may adopt alternative phenotypes. Taken together, our data supports the hypothesis that ST2 regulates the accumulation of effector T_regs_ in the tumor microenvironment over time by promoting the expression of DP signature genes.

### T_reg_-specific ST2 ablation leads to increased CD8+ T cell infiltration and a reduction in tumor burden

Finally, we found that tumors from KPfrt, *Foxp3*^*YFP-Cre*^, *Il1rl1*^*fl/fl*^ mice had over 50% higher CD8^+^ T cell infiltration than tumors from control mice by immunohistochemistry (**Figure 7E**). KPfrt, *Foxp3*^*YFP-Cre*^, *Il1rl1*^*fl/fl*^ mice also had a significantly lower total tumor burden and lower average tumor size compared to control mice (**Figure 7F,G**), suggesting that greater CD8+ T cell infiltration of tumors may result in better inhibition of tumor growth. Overall, our studies suggest that T_reg_-specific inhibition of ST2 signaling may result in a less immunosuppressive tumor microenvironment characterized by increased anti-tumor CD8 T cell activity and lower tumor burden.

## DISCUSSION

To identify specific features of T_regs_ in the tumor microenvironment that can be targeted therapeutically without adversely affecting T_regs_ in other tissues, we profiled T_conv_ and T_regs_ longitudinally in a mouse model of lung adenocarcinoma by scRNA-Seq. We show that T_reg_ diversity undergoes temporal shifts that would be missed in analyses of bulk populations or at a single timepoint. Leveraging these dynamic changes, we identified IL-33 as a critical mediator of effector T_reg_ function in tumors. Although previous scRNAseq studies have defined signatures of T_reg_ exhaustion and activation in human cancer (Guo et al., 2018; Zheng et al., 2017), our study is the first to effectively impair tumor growth by characterizing and perturbing a major pathway responsible for the development of transcriptionally-distinct subsets of T_regs_ in tumor-bearing lungs.

IL-33 has been shown to promote tumorigenesis through the recruitment of T_regs_ and other cells in transplant and xenograft models of breast and lung cancer (Jovanovic et al., 2014; Wang et al., 2016, 2017), and mice with T_reg_-specific ST2 deficiency have impaired growth of a transplantable tumor model (Magnuson et al., 2018). Here, we show in a genetically-defined, autochthonous mouse model of lung adenocarcinoma that loss of T_reg_-specific ST2 function is sufficient to impair tumor development without provoking systemic autoimmunity. Several therapeutic antibodies directed against ST2 and IL-33 are in preclinical development for the treatment of allergy and asthma. Our data point to the potential value of disrupting ST2 signaling in cancer.

Although ST2-deficient T_regs_ have been reported to be equally immunosuppressive as their wild-type counterparts *in vitro* (Schiering et al., 2014), our results suggest that *in vitro* suppression assays may fail to capture the full spectrum of T_reg_ effector activity *in vivo*. We observed a slight reduction in lung T_reg_ cell numbers as a result of T_reg_-specific ST2 deficiency, which may be related to reports that IL-33 can stimulate TCR-independent expansion of T_regs_ (Arpaia et al., 2015; Kolodin et al., 2015). DP T_regs_ from KPfrt, *Foxp3*^*YFP-Cre*^, *Il1rl1*^*fl/fl*^ mice had lower expression of Eff_TR genes compared to wild-type DP T_regs_, suggesting that ST2 may promote the maintenance of the effector T_reg_ phenotype. Indeed, IL-33 has been shown to increase expression of Foxp3 and GATA-3 (Kolodin et al., 2015; Vasanthakumar et al., 2015), transcription factors integral for T_reg_ terminal differentiation. Taken together, ST2-deficient T_regs_ may adopt an alternate functional state due to loss of IL-33 signaling.

T_regs_ across multiple tumor types likely have a common transcriptional program that is closely related to that of healthy tissue T_regs_ (Magnuson et al., 2018). Indeed, the effector T_regs_ in the KP model express a program similar to that of T_regs_ in several human cancers (De Simone et al., 2016; Guo et al., 2018), including a TNFRSF9^+^ T_reg_ population in human NSCLC (Zheng et al., 2017). This similarity may be due to the fact that clinically-detectable tumors are likely to have convergent strategies for evading immune destruction by recruiting highly suppressive T_regs_. Tumor-bearing lungs from KP mice also harbor Th1-like CXCR3^+^ T_regs_, which express IFN-stimulated genes and peak early in tumor development, following an opposite temporal pattern from the Eff_TR program. CXCR3 directs T_regs_ to sites of Th1 inflammation (Koch et al., 2009), which may explain the prominence of the IFNstim_TR program during early tumorigenesis, when CD8^+^ T cell infiltration of tumors and IFN signaling are most robust (DuPage et al., 2011). Cxcr3 may mark recently-arrived T_regs_ that have distinct functions from effector T_regs_, and temporal shifts in IFNstim_TR and Eff_TR gene expression may reflect T_reg_ adaptation to the tumor microenvironment over time. Alternatively, the decline in Cxcr3^+^ T_regs_ during tumor development may reflect cellular turnover and/or the outgrowth of an alternate subset of T_regs_ due to reduced IFN, and availability of IL-33 ligand. Several reports have described an IFN signature or a distinct population of CXCR3^+^ T_regs_ in human tumors, although their functional significance is not well-defined (Halim et al., 2017; Johdi et al., 2017; Redjimi et al., 2012).

Longitudinal profiling in the KP model provides a window into the natural history of effector T_reg_ activity that is challenging to achieve using patient samples. While Tr17-like, CXCR3+, and effector T_reg_ populations have been described previously in human tumors, we have shown that these states exist simultaneously, and their relative proportions vary with tumor development. Future studies may help elucidate the contribution of each distinct T_reg_ subset to tumor immune responses. While T_reg_ transcriptional heterogeneity may pose a challenge for efforts to target tumor T_reg_ activity, we show that loss of T_reg_-specific ST2 signaling can alter T_reg_ composition and ultimately impact tumor growth. Our study provides proof of concept that pathways that control T_reg_ diversity, maturation, and function may be useful targets for future therapies.

## EXPERIMENTAL METHODS

### Mice

KP, KPfrt, *Foxp3*^*GFP*^, *Foxp3*^*RFP*^, *Foxp3*^*GFP/DTR*^, *Il1rl1*^*−/−*^ and *Il1rl1*^*fl/fl*^ mice have been previously described (Bettelli et al., 2006; Chen et al., 2015; DuPage et al., 2011; Kim et al., 2007; Townsend et al., 2000; Wan and Flavell, 2005; Young et al., 2011). Both male and female mice were used for all experiments, and mice were gender and age-matched within experiments. Experimental and control mice were co-housed whenever appropriate. All studies were performed under an animal protocol approved by the Massachusetts Institute of Technology (MIT) Committee on Animal Care. Mice were assessed for morbidity according to MIT Division of Comparative Medicine guidelines and humanely sacrificed prior to natural expiration.

For *in vivo* labelling of circulating immune cells, anti-CD4-PE (eBioscience, RM4-4, 1:400) and anti-CD8β-PE (eBioscience, 1:400) were diluted in PBS and administered by IV injection 5 minutes before harvest (Anderson et al., 2012). Alternatively, anti-CD45-PE-CF594 (30-F11, BD Biosciences, 1:200) was also used for intravascular labeling and was administered 2 minutes before sacrifice.

For rIL-33 treatment studies, 200ng of recombinant mouse IL-33 (BioLegend) was diluted in 50 mL of PBS and administered intratracheally to mice as described previously (Li et al., 2014). Control mice received PBS only.

### Lentiviral production and tumor induction

The lentiviral backbone Lenti-LucOS has been described previously (DuPage et al., 2011). Lentiviral plasmids and packaging vectors were prepared using endo-free maxiprep kits (Qiagen). The pGK::GFP-LucOS::EFS::FlpO lentiviral plasmid was cloned using Gibson assembly (Akama-Garren et al., 2016; Gibson et al., 2009). Briefly, GFP-OS was created as a protein fusion of GFP and ovalbumin_257-383_, which includes the SIINFEKL and AAHAEINEA epitopes, and SIYRYYGL antigen. Lentiviral plasmids and packaging vectors were prepared using endo-free maxiprep kits (Qiagen). Lentiviruses were produced by co-transfection of 293FS* cells with Lenti-LucOS or FlpO-GFP-OS, psPAX2 (gag/pol), and VSV-G vectors at a 4:2:1 ratio with Mirus TransIT LT1 (Mirus Bio, LLC). Virus-containing supernatant was collected 48 and 72h after transfection and filtered through 0.45mm filters before concentration by ultracentrifugation (25,000 RPM for 2 hours with low decel). Virus was then resuspended in 1:1 Opti-MEM (Gibco) - HBSS. Aliquots of virus were stored at −80°C and titered using the GreenGo 3TZ cell line (Sánchez-Rivera et al., 2014).

For tumor induction, mice between 8-15 weeks of age received 2.5 × 10^4^ PFU of Lenti-LucOS or 4.5 × 10^4^ PFU of FlpO-GFP-OS intratracheally as described previously (DuPage et al., 2009).

### Tissue isolation and preparation of single cell suspensions

After sacrifice, lungs were placed in 2.5mL collagenase/DNAse buffer (Joshi et al., 2015) in gentleMACS C tubes (Miltenyi) and processed using program m_impTumor_01.01. Lungs were then incubated at 37°C for 30 minutes with gentle agitation. The tissue suspension was filtered through a 100 μm cell strainer and centrifuged at 1700 RPM for 10 minutes. Red blood cell lysis was performed by incubation with ACK Lysis Buffer (Life Technologies) for 3 minutes. Samples were filtered and centrifuged again, followed by resuspension in RPMI 1640 (VWR) supplemented with 1% heat-inactivated FBS and 1X penicillin-streptomycin (Gibco), and 1X L-glutamine (Gibco).

Spleens and lymph nodes were dissociated using the frosted ends of microscope slides into RPMI 1640 supplemented with 1% heat-inactivated FBS and 1X penicillin-streptomycin (Gibco), and 1X L-glutamine (Gibco). Spleen cell suspensions were spun down at 1500 RPM for 5 minutes, and red blood cell lysis with ACK Lysis Buffer was performed for 5 minutes. Cells were filtered through 40 μm nylon mesh and, after centrifugation, resuspended in supplemented RPMI 1640. Lymph node suspensions were filtered through a 40 μm nylon mesh, spun down at 1500 RPM for 5 minutes, and resuspended in supplemented RPMI 1640.

For *ex vivo* T cell stimulation experiments to detect intracellular cytokines, 0.5 × 10^5^ cells were plated in a 96-well U-bottom plate (BD Biosciences) in RPMI 1640 (VWR) supplemented with 10% heat-inactivated FBS, 1X penicillin-streptomycin (Gibco), 1X L-glutamine (Gibco), 1X HEPES (Gibco), 1X GlutaMAX (Gibco), 1mM sodium pyruvate (Thermo Fisher), 1X MEM non-essential amino acids (Sigma), 50μM beta-mercaptoethanol (Gibco), 1X Cell Stimulation Cocktail (eBioscience), 1X monensin (BioLegend), and 1X brefeldin A (BioLegend). Cells were incubated in a tissue culture incubator at 37°C with 5% CO_2_ for 4 hours.

### Staining for flow cytometric analysis

Approximately 0.5-1 × 10^6^ cells were stained for 15-30 minutes at 4°C in 96-well U-bottom plates (BD Biosciences) with directly conjugated antibodies (**Table S8**). SIINFEKL-Kb tetramer was prepared using streptavidin-APC (Prozyme) and SIINFEKL-Kb monomer from the NIH Tetramer Core.

After staining, cells were fixed with Cytofix/ Cytoperm Buffer (BD). Samples that were destined for Foxp3 or other transcription factor staining were fixed with the Foxp3 Transcription Factor Staining Buffer Kit (eBioscience). Intracellular cytokine and transcription factor staining were performed right before analysis using either the BD Perm/Wash Buffer (BD) or the Foxp3 Transcription Factor Staining Buffer Kit (eBioscience); staining was performed for 45 minutes at 4°C. Analysis was performed on an LSR II (BD) with 405, 488, 561, and 635 lasers. Data analysis was performed using FlowJo software.

### Isolation of T_reg_ populations for bulk RNA-Seq

For sequencing of LucOS-infected, KP, Foxp3-RFP mice: 100-200 DP, SP, and DN Treg cells were sorted into Buffer TCL (Qiagen) plus 1% b-mercaptoethanol using a MoFlo Astrios cell sorter. cDNA was prepared by the SMART-Seq2 protocol (Picelli et al., 2013) with the following modifications: RNA was purified using 2.2X RNAclean SPRI beads (Beckman Coulter) without final elution, after which beads were air-dried and immediately resuspended with water and oligoDT for annealing, and 18 cycles of preamplification were used for cDNA. cDNA was then mechanically sheared and prepared into sequencing libraries using the Thru-Plex-FD Kit (Rubicon Genomics). Sequencing was performed on an Illumina HiSeq 2000 instrument to obtain 50 nt paired-end reads.

For comparison of wild-type and ST2-deficient Tregs and CD8+ T cells: 100-200 DP, SP, and DN Tregs or SIINFEKL-tetramer-positive and negative CD8^+^ T cells were sorted and cDNA was prepared with 14 cycles of preamplification. Nextera library preparation was performed as previously described (Picelli et al., 2013) and sequencing was performed with 50 × 25 paired end reads using two kits on the NextSeq500 5 instrument.

### Single-cell sorting of T_conv_ and T_reg_ populations for RNA sequencing

Tconv (DAPIneg, i.v. neg, Thy1.2+CD4+Foxp3-GFPneg) and Treg (DAPIneg, i.v. neg, Thy1.2+CD4+Foxp3-GFP-positive) cells were single-cell sorted into Buffer TCL (Qiagen) plus 1% B-mercaptoethanol in 96-well plates using a MoFlo Astrios cell sorter. Each plate had 30-100 cell population well and an empty well as controls. Following sorting, plates were spun down for 1” at 2000 RPM and frozen immediately at −80C.

### Preparation of scRNAseq libraries

Plates were thawed and RNA was purified using 2.2X RNAclean SPRI beads (Beckman Coulter) without final elution (Shalek et al., 2013). SMART-seq2 and Nextera library preparation was performed as previously described (Picelli et al., 2013), with some modifications as described in a previous study (Singer et al., 2017). Plates were pooled into 384 single-cell libraries, and sequenced 50 × 25 paired end reads using a single kit on the NextSeq500 5 instrument.

### Quantitative PCR for validation of RNA-Seq experiments

Quantitative PCR was performed using various primer sets (**Table S5**). 1ng of cDNA generated using SMART-Seq2 was included in a reaction with 1μL of each primer (2μM stock) and 5μL of KAPA SYBR Fast LightCycler 480 (KAPA Biosystems). Cp values were measured using a LightCycler 480 Real-Time PCR System (Roche). Relative fold-change in expression values were calculated using the following formula: 2^(ΔCp(Sample) - ΔCp(Spleen))^, where (ΔCp(Sample) = Sample Cp_Gene of Interest_ - Sample Cp_GAPDH_, and ΔCp(Spleen) = Spleen Cp_Gene of Interest_ - Spleen Cp_GAPDH_.

### Population-level TCR Beta chain sequencing and analysis

For bulk TCR beta chain sequencing, T cells were sorted directly into 250μl RNAprotect buffer (Qiagen), spun down for 1 minute at 2000 RPM, and immediately frozen at −80°C. Samples were sent to iRepertoire (Huntsville, AL) for library preparation and sequencing. TCR sequences were analyzed and compared with VDJtools software (Shugay et al., 2015).

### Immunohistochemistry (IHC) and immunofluorescence staining

Lung lobes and spleens allocated for IHC and IF were perfused with 4% paraformaldehyde in PBS and fixed overnight at 4°C. Lung lobes and/ or spleen were transferred to histology cassettes and stored in 70% ethanol until paraffin embedding and sectioning (KI Histology Facility). H&E stains were performed by the core facility using standard methods.

For IHC, 5 μm unstained slides were dewaxed, boiled in citrate buffer (1 g NaOH, 2.1 g citric acid in 1L H2O, pH 6), for 5 minutes at 125°C in a decloaking chamber (Biocare Medical), washed with 3X with 0.1% Tween-20 (Sigma) in TBS, and blocked and stained in Sequenza slide racks (Thermo Fisher). Slides were blocked with Dual Endogenous Peroxidase and Alkaline Phosphatase Block (Dako) and then with 2.5% Horse Serum (Vector Labs). Slides were incubated in primary antibody overnight, following by washing and incubation in HRP-polymer-conjugated secondary antibodies (ImmPRESS HRP mouse-adsorbed anti-rat and anti-goat, Vector Laboratories). Slides were developed with ImmPACT DAB (Vector Laboratories). Primary antibodies used were goat anti-IL-33 (R&D, AF3626) and rat anti-CD8a (Thermo Fisher, 4SM16). Stains were counterstained with hematoxylin using standard methods before dehydrating and mounting.

After fixation, lung lobes and spleen allocated for IF were perfused with 30% sucrose in PBS for cryoprotection for 6-8h at 4°C. Tissues were then perfused with 30% optimum cutting temperature (O.C.T.) compound (Tissue-Tek) in PBS and frozen in 100% O.C.T in cryomolds on dry ice. 6μm sections were cut using a CryoStar NX70 cryostat (Thermo), and air-dried for 60-90 minutes at room temperature. Sections were incubated in ice-cold acetone (Sigma) for 10 minutes at −20°C and then washed 3 x 5 minutes with PBS. Samples were permeabilized with 0.1% Triton-X-100 (Sigma) in PBS followed by blocking with 0.5% PNB in PBS (Perkin Elmer). Primary antibodies were incubated overnight. Primary antibodies used were rabbit anti-prosurfactant protein C (SPC) (Millipore, AB3786, 1:500) and goat anti-IL-33 (R&D, AF3626, 1:200). After washing 3 x 5 minutes, samples were incubated in species-specific secondary antibodies conjugated to Alexa Fluor 568 and Alexa Fluor 488, respectively, at 1:500. Sections were then fixed in 1% PFA and mounted using Vectashield mounting media with DAPI (Vector Laboratories).

Immunohistochemistry and immunofluorescence tissue section images were acquired using a Nikon 80 Eclipse 80i fluorescence microscope using 10x and 20x objectives and an attached Andor camera. Stained IHC slides were scanned using the Aperio ScanScope AT2 at 20X magnification.

## COMPUTATIONAL ANALYSIS

### Bulk RNA-seq data processing and signature analyses

Bulk RNA-Seq reads that passed quality metrics were mapped to the annotated UCSC mm9 mouse genome build (http://genome.ucsc.edu/) using RSEM (v1.2.12) (http://deweylab.github.io/RSEM/) (Li and Dewey, 2011) using RSEM’s default Bowtie (v1.0.1) alignment program (Langmead et al., 2009). Expected read counts estimated from RSEM were upper-quartile normalized to a count of 1000 per sample(Bullard et al., 2010). Genes with normalized counts less than an upper-quartile threshold of 20 across all samples were considered lowly expressed and excluded from further analyses. The dataset was log_2_ transformed before subsequent analysis.

Unsupervised clustering of samples was performed using a Pearson correlation-based pairwise distance measure.

Signature analyses between bulk Treg cell populations were performed using a blind source separation methodology based on ICA (Hyvärinen and Oja, 2000), using the R implementation of the core JADE algorithm (Joint Approximate Diagonalization of Eigenmatrices)(Biton et al., 2014; Nordhausen et al., 2014; Rutledge and Jouan-Rimbaud Bouveresse, 2013) along with custom R utilities. Multi-sample signatures were visualized using relative signature profile boxplots (Li et al., 2018). Signature correlation scores (z-scores) for each gene are included in Tables S1 and S7. Heat maps were generated using the Heatplus package in R.

### Gene Set Enrichment Analysis (GSEA)

Gene set enrichment analyses were carried out using the pre-ranked mode in GSEA with standardized signature correlation scores for the KPLungTR signature and default settings using gene-sets from MsigDB v5.1 (Subramanian et al., 2005) and a custom immunologic signatures library of gene sets (**Table S2**) added to version 4.0 of the MSigDB immunologic collection (c7). Normalized Enrichment Score (NES), p-values and FDR for the custom gene-sets were calculated in the context of the combined c7 v4.0 MSigDB collection.

Network representations of GSEA results were generated using EnrichmentMap (http://www.baderlab.org/Software/EnrichmentMap) for Cytoscape v3.3.0 (http://www.cytoscape.org).

### Identification of DP signature

To identify a signature separating CD103^+^KLRG1^+^ lung T_regs_ from other populations we applied ICA to the data prior to log transformation, which allowed us to detect signatures with lower amplitudes of gene expression changes. We detected a signature separating CD103^+^KLRG1^+^ lung Tregs from other populations. Genes in this signature with |z-score| > 3 were selected for downstream analysis (75 up-regulated and 31 down-regulated genes). An additional expression level filter was implemented to narrow the list of genes of interest. For upregulated genes, expression in all CD103^+^KLRG1^+^ lung Treg samples had to be greater than all but a maximum of 3 other samples (3 out of a total 8 other samples). A similar filtering scheme was employed in the other direction for down-regulated genes. This yielded a total of 43 up-regulated and 2 down-regulated genes in CD103^+^KLRG1^+^ lung Tregs (**Table S1**). This set of genes was used to illustrate gene expression level changes in a heatmap (**Figure 1D**).

### Filtering of genes differentially-expressed in ST2-deficient T_regs_

A signature distinguishing ST2-deficient Tregs from wild-type Tregs was identified through ICA (**Table S7**). To identify particular genes of interest, signature genes (|z-score| > 3) were filtered to include only genes that had an absolute fold change exceeding 1.5x within any of the CD103^+^KLRG1^+^ (DP), CD103^+^KLRG1^−^(SP), CD103^−^KLRG1^−^(DN) sample types between wild-type and ST2-deficient Tregs. These gene lists were then filtered to retain only those genes that appeared in at least two of the three sample types (i.e. up/down-regulated in wild-type or ST2-deficient in at least two of DP/DN/SP comparisons). Genes with opposite directionality across the three sample types (n=5 genes) were dropped. Expression levels of the resulting curated set of 14 genes were visualized using a row-normalized heatmap (**Figure 7D**).

### Pre-processing of SMART-Seq2 scRNA-seq data

BAM files were converted to de-multiplexed FASTQs using the Illumina-provided *Bcl2Fastq* software package v2.17.1.14. Paired-end reads were mapped to the UCSC mm10 mouse transcriptome using Bowtie with parameters ‘-n 0 -m 10’, which allows alignment of sequences with zero mismatches and allows for multi-mapping of a maximum of ten times.

Expression levels of genes were quantified using TPM values calculated by RSEM v1.2.8 in paired-end mode. For each cell, the number of detected genes (TPM > 0) was calculated and cells with less than 600 or more than 4,000 genes detected were excluded as well as cells that had a mapping rate to the transcriptome below 15%. To further remove potential doublets (mostly of B cells and epithelial cells), we calculated the sum log_2_(TPM+1) over *Cd79a*, *Cd19*, *Lyz1*, *Lyz2* and *Sftpc*, and excluded any cell that scored higher than 3. We retained only genes expressed above log_2_TPM of 3 in at least five cells in the whole dataset.

Since we could not sort for T_reg_ for two of the mice (#336 and #338), we had to infer which cells are T_regs_ from these data. To this end, we trained a random forest classifier for mice for which we have sorted both T_conv_ and T_regs_, using the train function from the *caret* package in R, based on the expression of the following genes: *Foxp3*, *Ikzf2*, *Areg*, *Il1rl1*, *Folr4*, *Wls*, *Tnfrsf9*, *Klrg1*, *Il2ra*, *Dusp4*, *Ctla4*, *Neb*, *Itgb1*, and *Cd40lg*. The labeled data was partitioned into training and test sets. The model has a sensitivity and specificity above 90% in cross validation. We then applied the classifier on the unlabeled data and cells with a probability above 0.6 to be either T_conv_ or T_reg_ were given the corresponding label. The remaining 4% of cells were discarded as unambiguous.

### Identifying tissue-specific gene programs for T_reg_ and T_conv_

To identify genes that are differentially expressed between lung and msLN in T_reg_ and/or T_conv_, we performed a regression analysis. We focused on the proportion of cells expressing a gene, and hence on logistic regression. We performed logistic regression using the bayesglm function from the *arm* package in R, including only those mice (# 338, #3642, #3839, #3889) for which we had matched cells from both lung and msLN, as well as for T_reg_ and T_conv_, and excluding all genes expressed in >95% or <5% of cells in lung and msLN. We ran the logistic regression with expression data binarized at a log_2_(TPM+1) of 2 and using the following full model: *gene expression ~ genes detected + batch effect + tissue* versus a reduced model: *gene expression ~ genes detected + batch effect*. We corrected for multiple hypothesis by computing an FDR of the likelihood ratio test p-value, and retained genes as differentially expressed between lung and msLN with P< 10^−5^ and an |coefficient| > 2.

### Comparing the extent of cell heterogeneity between lung and msLN

Diffusion components were calculated on a gene expression matrix limited to genes that were differentially expressed between lung and msLN using the DiffusionMap function from the *destiny* package in R (Angerer et al., 2016) with a *k* of 30 and a local sigma. In order to be able to compare the variance in distributions in diffusion component 1 and 2 between lung and msLN T_reg_/T_conv_, we downsampled the cells from the lung to the (lower) numbers of cells from the msLN. To test for significant differences in variance in the distributions of lung and msLN T_reg_/T_conv_, we used Levene’s test for the equality of variances on the distributions of the coefficients of the downsampled cells in each of diffusion components 1 and 2.

### Identifying gene modules and their time dependence

Gene modules were identified using PAGODA using the *scde* R package version 2.6.0. (Fan et al., 2016) on the counts table from RSEM after cleaning the data using the clean.counts function (min.lib.size=600,min.detected=5). The *knn.error.model* function was run using a *k* of 30, which is much lower than default, but yields statistically indistinguishable results from the default k (# cells / 4). We then ran the *pagoda.varnorm* to normalize gene expression variance, and the *pagoda.subtract.aspect* function to control for sequencing depth which then allowed us to run *pagoda.gene.clusters* which identifies *de-novo* correlated genes in the dataset. We forced PAGODA to return 100 modules. We identified modules with a significance z.score above 1.96. We removed several highly significant newly identified gene modules consisting of paralog groups with high expression correlation, likely because of multimapping of reads.

Mean module expression was calculated by averaging over the genes in each module of the centered and scaled gene expression table and transforming to a z-score over 1,000 randomly selected gene sets with matched mean-variance patterns. As an initial step, all genes were binned into 10 bins based on their mean expression across cells, and into 10 (separate) bins based on their variance of expression across cells. Given a gene signature (e.g. list of genes in a module), a cell-specific signature score was computed for each cell as follows: First, 1,000 random gene lists were generated, where each instance of a random gene-list was generated by sampling (with replacement) for each gene in the gene-list a gene that is equivalent to it with respect to the mean and variance bins it was placed in. Then, the sum of gene expression in the given cell was computed for all gene-lists (given the 1,000 random lists generated) and the z-score of the original gene-list for the generated 1,000 sample distribution is returned, as in (Singer et al., 2017).

Another module of highly correlated genes identified by PAGODA showed no biological relevance based on gene annotation, but was associated with cells processed on specific dates, suggested they reflect a contamination or batch effect. We scored each cell for this module with the above described method for scoring cells for gene signatures. When testing for differential gene expression over tumor development (described below), we included this batch effect score as a covariate in the regression analysis to control for genes that are correlated with it.

To test if a module’s expression changes over the course of tumor development, we estimated a linear model for each module and compared with a likelihood ratio test a full model: *module.activity ~ detected genes + time point* to a reduced model: *module.activity ~ detected genes.* For the time point covariate, healthy lung was taken as reference. We corrected the likelihood ratio test p-values for multiple hypotheses for the number of modules using the *p.adjust* function computing the false discovery rate in the *stats* package.

### Dimensionality reduction using diffusion component analysis

Diffusion components were calculated on a gene expression matrix limited to genes from modules of interest: modules 1,4,5,14,15 and 21 for T_conv_, and modules 1,3,6,8,9,12,13,18,21,23 and 26 for T_reg_. Gene expression was scaled for T_regs_ only across all cells. Diffusion components were calculated using the DiffusionMap function from the *destiny* package in R (Angerer et al., 2016) with a *k* of 30 and a local sigma. Significant diffusion components identified by the elbow in the eigenvalues were further used for dimensionality reduction to two dimensions. The eigenvectors of the significant diffusion components were imported into gephi 0.9.2 and a force directed layout using forceatlas 2 was run until it converged to get a two dimensional embedding.

### Testing for differential gene expression during tumor development

To test whether individual genes change in gene expression over the course of tumor growth, we performed a two-step regression analysis. We focused on the proportion of cells expressing a gene, and hence on logistic regression. We performed logistic regression using the bayesglm function from the *arm* package in R. Because gender is often confounded with a particular time point in our experiment, we did not include it as a covariate in the model, but did remove all Y chromosome genes from analysis. We also excluded all genes expressed in >95% or <5% of cells in each mouse. We ran the logistic regression with expression data binarized at a log2(TPM+1) of 2 and using the following full model: *gene expression ~ genes detected + batch effect + week p.i.* (healthy lung as reference) versus a reduced model: *gene expression ~ genes detected + batch effect*. We identified a threshold for significance by the elbow method, identifying the peak of the second derivative of the ordered fdr distribution of the likelihood ratio test for each time point. To remove significant genes whose signal was driven by only one mouse, we performed another logistic regression using a mixed effect model, accounting for mouse variability: To this end, we added to the significant genes 1,000 randomly selected genes that were non-significant by the initial test to serve as background genes, and performed a mixed effect logistic regression using the glmer function of the lme4 package in R, with the model *gene expression ~ tmp + (1|mouse)*, allowing the intercept to vary by mouse. We combined the elbow method above and the background genes to select an FDR cutoff for significance of 0.01. A gene was classified as significantly varying during tumor development if it passed this FDR cutoff in at least one time point.

### T cell receptor (TCR) reconstruction and clonotype calling

TCR were reconstructed using Tracer (Stubbington et al., 2016), run in short read mode with the following settings ‘--inchworm_only=T --trinity_kmer_length=17’. To call shared clonotypes between T_reg_ and T_conv_ cells, we required all cells of a clone to have identical productive TCRA and TCRB.

### Comparison of bulk and scRNA-seq signatures to published signatures

Lists of differentially expressed genes in human cancer T_regs_, mouse tissue T_regs_, Tr17 cells from mice, and mouse activated T_regs_ (**Table S4**) were collected either from the supplementary tables of the relevant publications, or generously provided by the authors upon request (De Simone et al., 2016; Kim et al., 2017; Miragaia et al., 2017; Plitas et al., 2016; van der Veeken et al., 2016).

### Population-level TCR Beta chain sequencing and analysis

#### Analysis of IHC Images

QuPath software was used to annotate tumor and lobe areas (Bankhead et al., 2017). CD8-stained images were standardized to a common set of stain vector parameters. CD8^+^ cell detection was performed using the PositiveCellDetection plugin with the following parameters: runPlugin(‘qupath.imagej.detect.nuclei.PositiveCellDetection’, ‘{“detectionImageBrightfield”: “Optical density sum”, “requestedPixelSizeMicrons”: 0.5, “backgroundRadiusMicrons”: 8.0, “medianRadiusMicrons”: 0.0, “sigmaMicrons”: 1.5, “minAreaMicrons”: 7.0,“maxAreaMicrons”: 125.0, “threshold”: 0.3, “maxBackground”: 2.0, “watershedPostProcess”: true, “excludeDAB”: false, “cellExpansionMicrons”: 2.0, “includeNuclei”: false,“smoothBoundaries”: false, “makeMeasurements”: true, “thresholdCompartment”: “Cytoplasm: DAB OD max”, “thresholdPositive1”: 0.7, “thresholdPositive2”: 0.4, “thresholdPositive3”: 0.6, “singleThreshold”: true}’);

Scored cells were normalized to tumor area.

### Additional statistical analyses

Unpaired, two-tailed Student’s t tests, Mann-Whitney tests, Tukey’s multiple comparisons tests, and Sidak’s multiple comparisons tests were used for all statistical comparisons using GraphPad Prism software.

## AUTHOR CONTRIBUTIONS

A.L., R.H.H., D.C., J.M.S., C.G.R., M.B., L.C., A.R., and T.J. designed the study; A.L., D.C., and J.M.S. performed all of the mouse experiments and collection of samples for RNA-Seq in the laboratory of T.E.J.; R.H. performed all computational analysis of RNA-Seq data in the lab of A.R., with help from A.B.; C.D. and M.H. provided technical assistance; C.G.R. conducted TCR repertoire analyses in the laboratory of M.B.; L.C., O.C.S., J.Y.K., and M.C. performed scRNA-seq in the laboratory of A.R., under guidance and supervision from O.R.R.; P.S.R. assisted with cell sorting. A.L., R.H.H., D.C., J.M.S., A.R., and T.J. wrote the manuscript with input from other authors.

## ACKNOWLEDGEMENTS

We thank N. Joshi, N. Marjanovic, R. Satija, D. Gennert, K. Thai, and C. Jin for thoughtful discussions,and technical advice; S. Riesenfeld for computational advice; J. Park, J. Wilson and N. Cheng for technical assistance with animal experiments; S. Levine at the MIT BioMicro Center for sequencing support; C. Otis and S. Saldi in the Broad Flow Cytometry Core for sorting assistance; G. Paradis in the Koch Institute Flow Cytometry Facility for technical advice on flow cytometry; K. Cormier and C. Condon from the Hope Babette Tang (1983) Histology Facility for histology assistance; L. Gaffney for artwork and advice on figures; A. Rudensky for *Foxp3*^*DTR-GFP*^ mice; A. Sharpe for *Foxp3*^*GFP*^ mice; D. Artis for *Il1rl1*^−/−^ mice; D. Mathis for *Il1rl1*^*fl/fl*^ mice; K. Anderson, J. Teixeira, M. Magendantz, and K. Yee for administrative and logistical support.

This work was supported by the Howard Hughes Medical Institute (T.J. and A.R.), Margaret A. Cunningham Immune Mechanisms in Cancer Research Fellowship Award (A.L.), David H. Koch Graduate Fellowship Fund (A.L.), NCI Cancer Center Support Grant P30-CA1405, an Advanced Medical Research Foundation grant (D.C.), and by the Klarman Cell Observatory at the Broad Institute. A.L. is supported by T32GM007753 from the National Institute of General Medical Sciences. A.R. is an Investigator of the Howard Hughes Medical Institute, SAB member for Thermo Fisher and Syros Pharmaceuticals, and a consultant for Driver Group. T.J. is a Howard Hughes Medical Institute Investigator, David H. Koch Professor of Biology, and a Daniel K. Ludwig Scholar.

## DISCLOSURES

T.J. receives research support from the J&J Lung Cancer Initiative. T.J. is a member of the Board of Directors of Amgen and Thermo Fisher Scientific and an equity holder in both companies. He is co-Founder and Scientific Advisory Board member of Dragonfly Therapeutics, a co-founder of T2 Biosystems, and a Scientific Advisory Board member of SQZ Biotech; he is an equity holder in all three companies. His laboratory currently receives funding from the Johnson & Johnson Lung Cancer Initiative and Calico.

---

**Figure S1, related to Figure 1.**
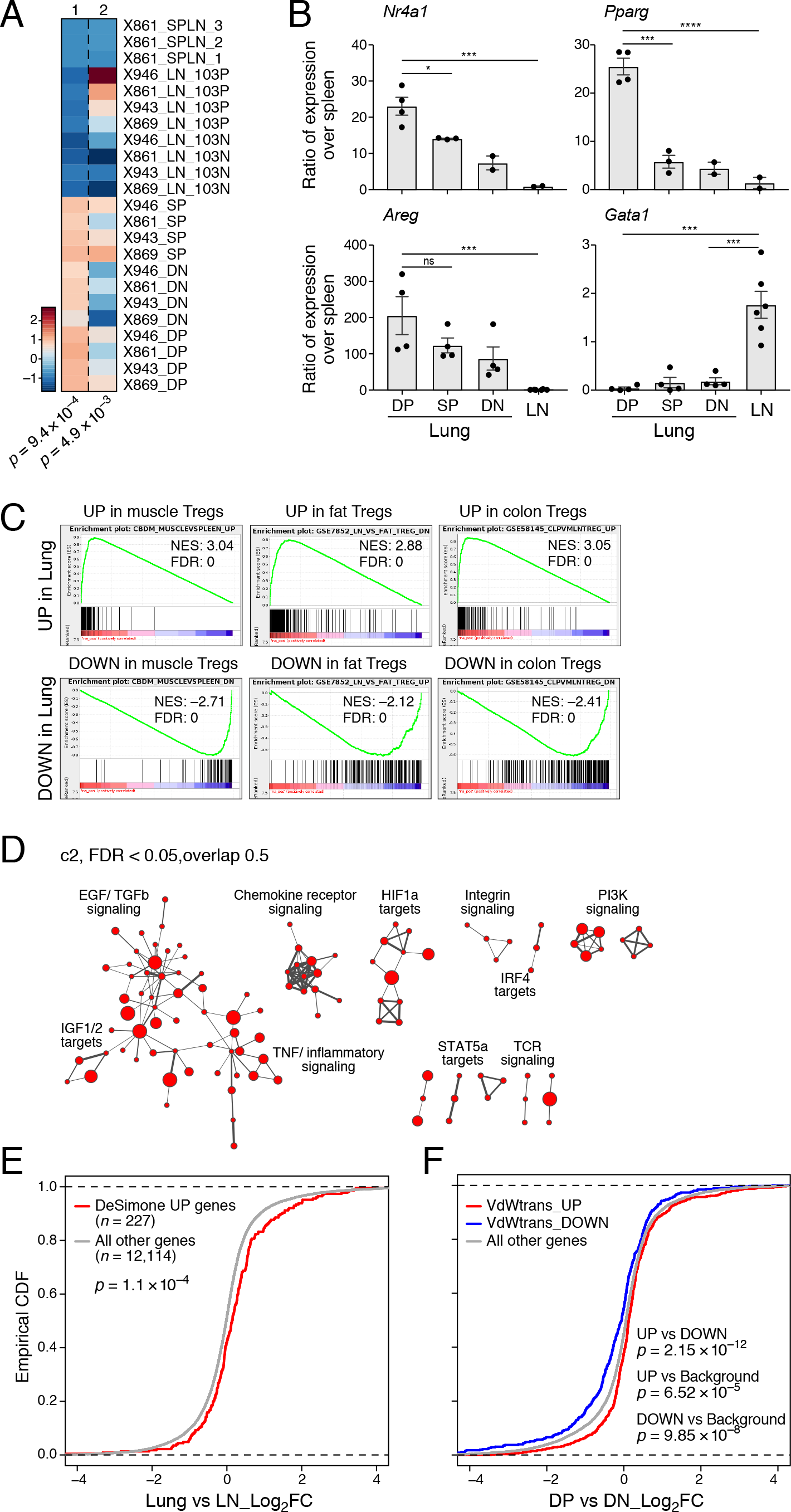
Characteristics of effector lung T_regs_ from tumor-bearing KP mice. **A.** Significant expression signatures identified by ICA. Mixing weight z-scores (color bar) per sample (row) for two gene expression signatures (columns). Signature 1 distinguishes lung populations (DP, DN, SP) from spleen and LN ones. Signature 2 distinguishes CD103^+^ T_regs_ from CD103^−^ populations. P-values for these distinctions: Kruskal-Wallis test. **B.** Validation of expression differences. qRT-PCR of expression of *Pparg*, *Nr4a1*, *Gata1*, and *Areg1* (y-axis, 2^ΔΔCt^ values, with splenic Treg expression as control) in DP, SP, and DN lung T_regs_ and in SP and DN msLN T_reg_ cells. Error bars: SEM. *p<0.05, ***p<0.001, ****p<0.0001, Tukey’s multiple comparisons test. NS: non-significant. **C-D.** GSEA of enriched functional categories in the KPLungTR signature. **C.** Test details for gene sets induced (top) or repressed (bottom) in the KPLungTR signature (**Methods**). **D.** Network representation of GSEA gene sets (nodes) from the curated collection (c2) enriched in the KPLungTR signature (p < 0.05, FDR < 0.05; in all significant gene sets, the upregulated genes were enriched). Node size: gene set size. Edge thickness: overlap between gene sets (minimum: 50% overlap). Related pathways were manually annotated. **E.** Signature enrichment for orthologs of genes included in human CRC and NSCLC-associated T_regs_. Empirical cumulative distribution functions (ECDFs) of Lung vs LN log_2_(fold-change) of expression for genes upregulated in CRC and NSCLC T_regs_ (DeSimone_UP, red) (De Simone et al., 2016) and all other expressed genes (gray). p = 1.137 ×10^4^, two-sided Kolmogorov – Smirnov test. **F.** DP cells have features similar to activated Tregs. ECDFs of DP vs. DN T_regs_ log_2_(fold-change) of expression of genes transiently upregulated (VdWtrans_UP,red), downregulated (VdWtrans_DN, blue) in activated T_reg_ cells (van der Veeken et al., 2016), or all other genes (gray). P-values: two-sided Kolmogorov – Smirnov test.

**Figure S2, related to Figure 2.**
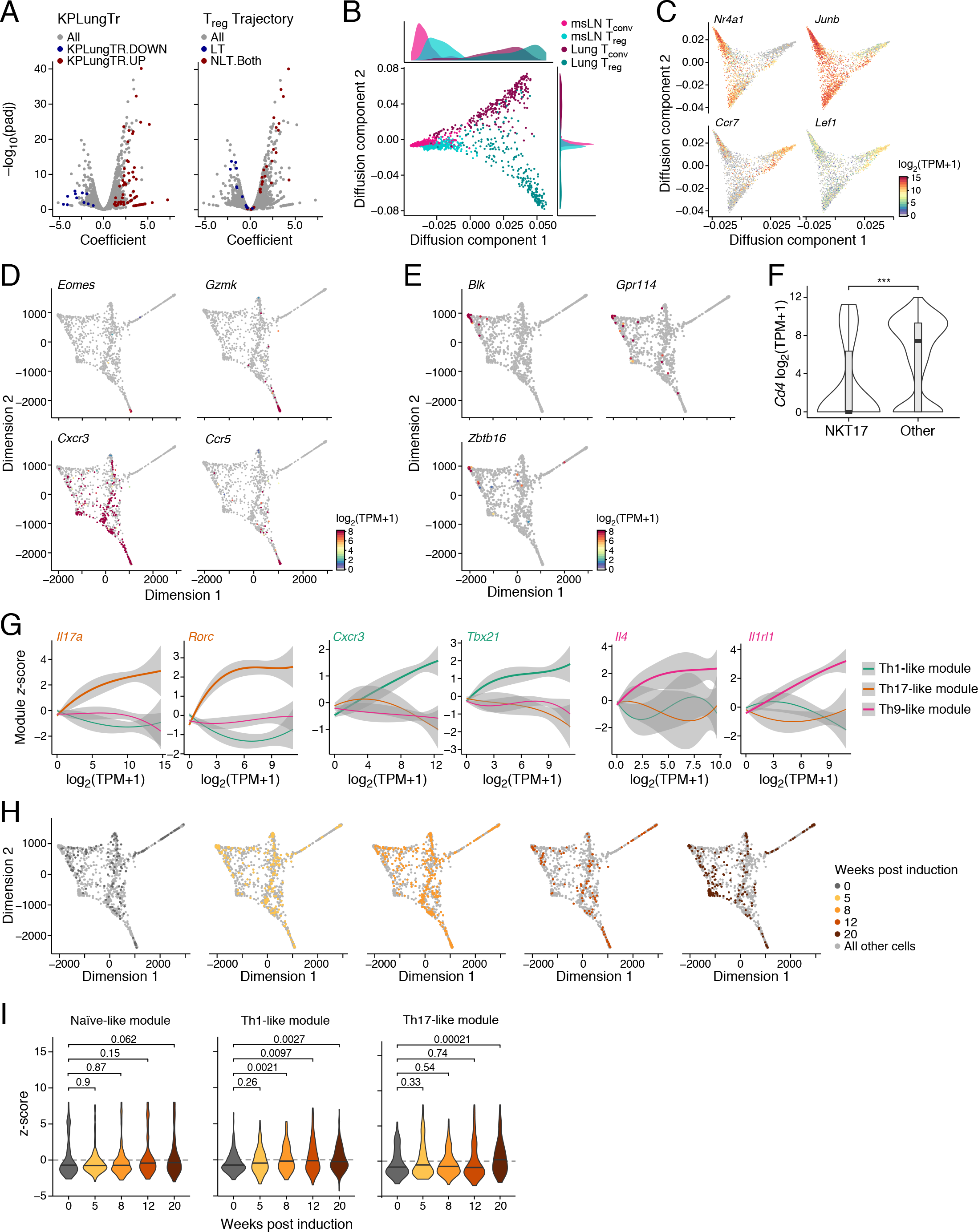
scRNAseq reveals lung CD4+ T_conv_ diversity in KP tumors. **A.** Differentially expressed genes between lung and msLN T_regs_. Shown for each gene (dot) is its differential expression between lung and msLN T_regs_ (x-axis) and associated significance on the y-axis, log_10_(p-value) (logistic regression, **Methods**). Red/blue genes are upregulated/ downregulated from the KPLungTR signature (left) or upregulated in both skin and colon compared to lymph node (Miragaia et al., 2017) (right), highlighting overlapping genes. **B.** Lung cells are more variable. Map of the first two diffusion components of T_reg_ and T_conv_ cells from the lung and msLN, where lung samples were downsampled to equal numbers as in msLN. Histograms: distribution of the cell scores in each diffusion component. **C.** Naive and central memory gene expression in CD4^+^ T cells. Diffusion component embedding for all CD4+ T cells (as in Figure 2C) colored by log2(TPM+1) expression (color bar) of *Ccr7* and *Lef1* (naive and central memory markers), and *Junb* and *Nr4a1* (T cell activation markers). **D-E.** Cytotoxic, Th1, and NKT17 cell-associated gene expression in T_conv_ lung cells. Two dimensional (2D) force directed layout embedding of T_conv_ lung cells (as in **Figure 2D**) colored by log2(TPM+1) expression (color bar) of *Eomes*, *Gzmk*, *Cxcr3* or *Ccr5* (D, cytotoxic and Th1 cells) or *Blk*, *Gpr114* and *Zbtb16* (E, NKT17 cells). **F.** *Cd4* is significantly downregulated in NKT17-like cells. Distribution of log_2_(TPM+1) *Cd4* gene expression (y-axis) of NKT17 cells and all other T_conv_ of the lung. p < 0.001, Kolmogorov-Smirnov test. **G.** Th1, Th17, and Th9 modules. Smoothed loess distribution of log_2_(TPM+1) expression (x-axis) of key genes (label top, color code) for the Th1 (green), Th17 (orange), and Th9 (red) modules in cells and the associated activity z-score (y-axis) of each module in these cells. Bold curve: score for module in which each gene is a member. **H.** Temporal changes. Two-dimensional force-directed layout embedding of T_conv_ lung cells (as in **Figure 2D**) colored by timepoint after tumor induction. **I.**T_conv_ subsets remain largely stable over tumor development. Distributions of module activity z-scores (y-axis) for each module (label, top). P-values: Kolmogorov-Smirnov test (vs. non tumor-bearing lung).

**Figure S3, related to Figure 3.**
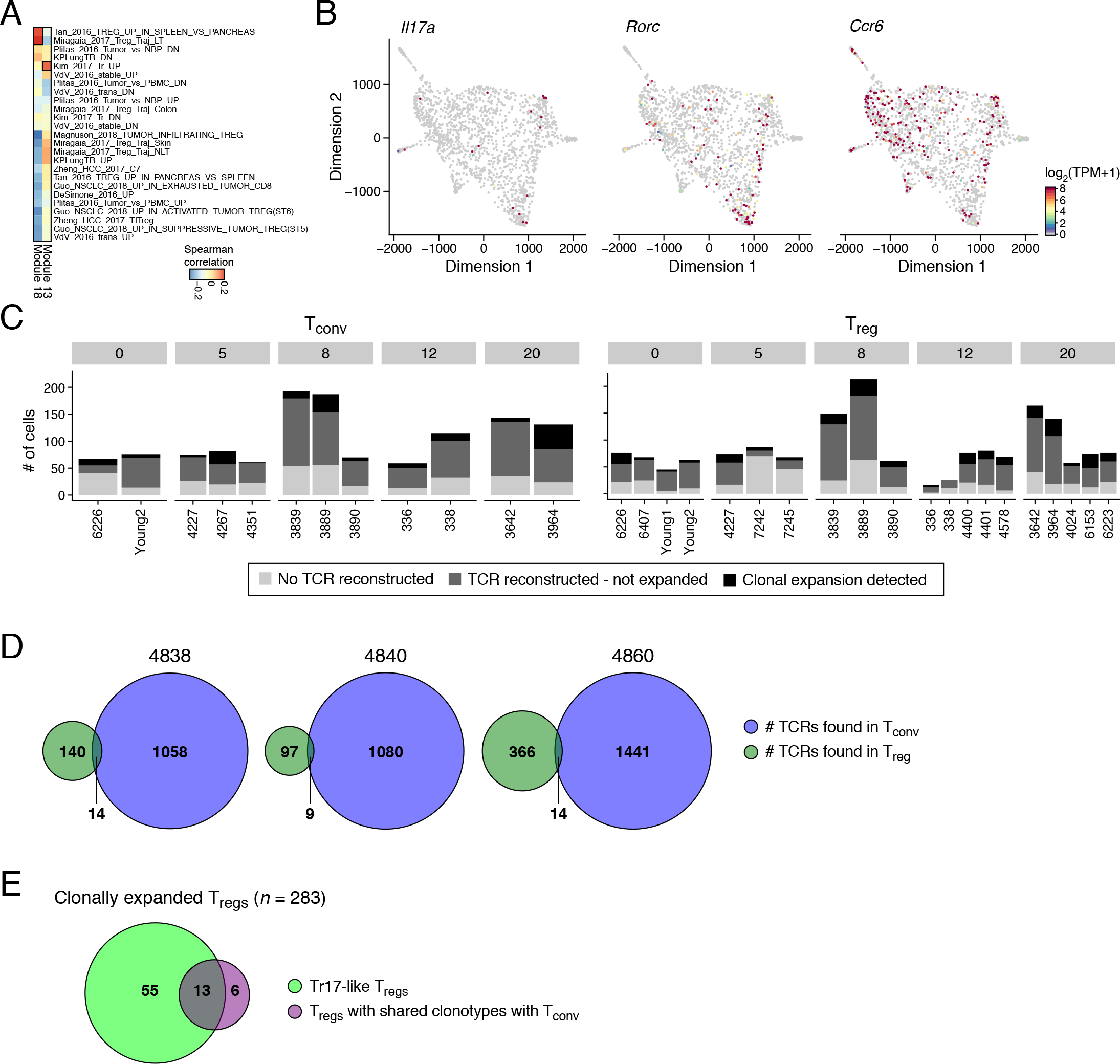
Th17-like T_reg_ population in tumor development. **A.** T_reg_ modules are associated with previously-described gene expression programs. Spearman correlation coefficient (color bar) between module (columns) z-scores across cells and z-scores for published signatures (rows, **Methods**, **Table S6**) of various T_reg_ states. **B.** Expression of key Th17 cell-associated genes. Two dimensional force directed layout embedding of T_reg_ lung cells (as in Figure 3C) colored by normalized expression (z-score) of *Il17a, Rorc* or *Ccr6*. **C.** T cell clones inferred by TCR reconstruction. Number of T_conv_ (left) and T_reg_ (right) cells (y-axis) in each mouse (x-axis) for which we did not identify a TCR (light gray), identified a TCR but not a shared clone (medium gray), or identified a clone (dark gray). **D.**Validation of shared clonotypes between T_reg_ and T_conv_ cells. Bulk TCR sequencing results of three replicates, showing the number of identified clonotypes in each subset and overall, and the overlap. We estimated that about 5% of T_reg_ clonotypes are shared with T_conv_. **E.** T_regs_ that have a shared clonotype with T_conv_ are enriched for Tr17-like cell. Numbers of Tr17-like cells (green), of T_regs_ with shared clonotype with T_conv_ (purple), and the overlap. p < 10^−5^, hypergeometric test.

**Figure S4, related to Figure 4.**
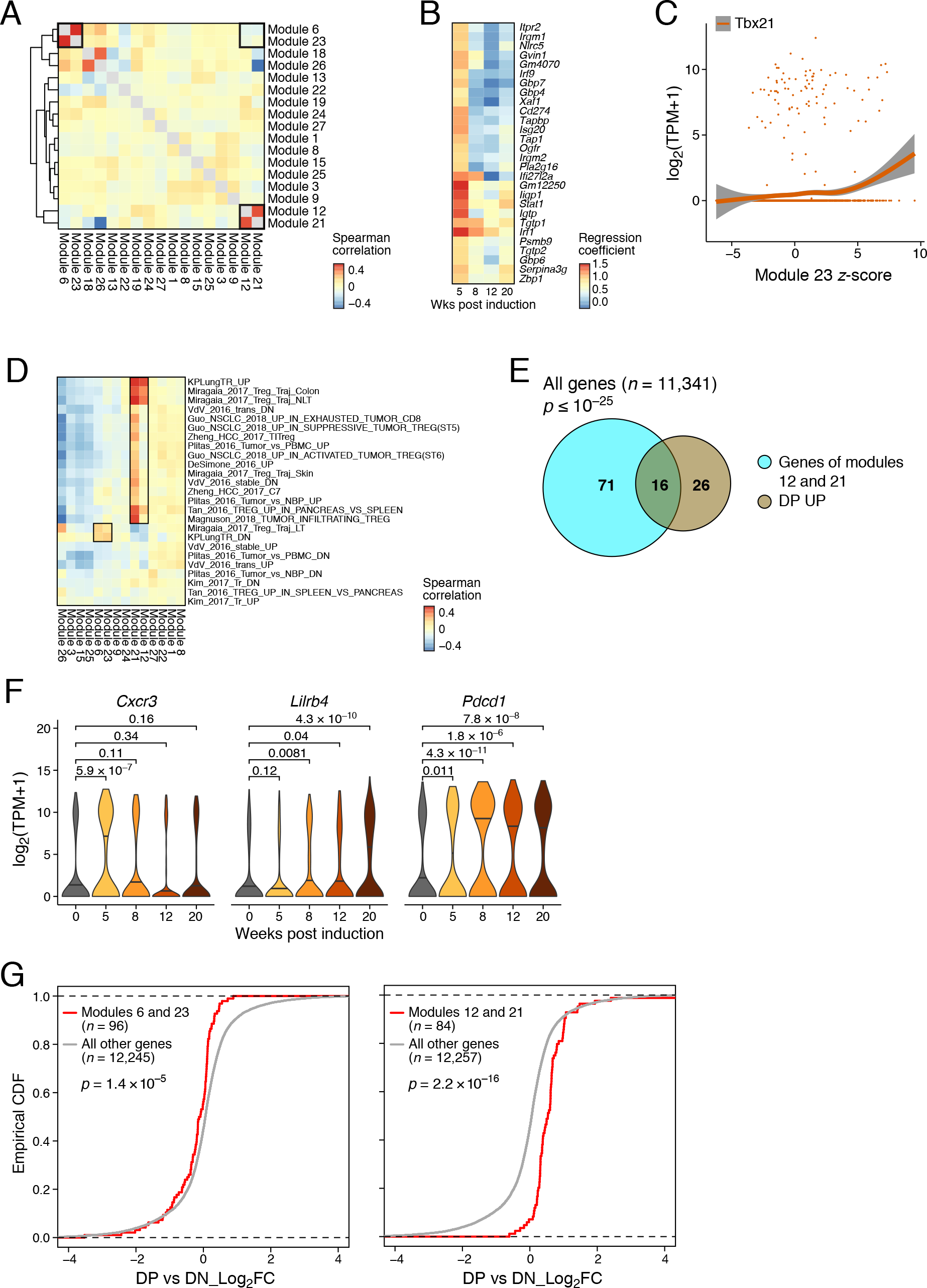
Effector Treg cells become predominant later in tumor development. **A.** Different modules pick up on similar signals and are correlated in expression across cells. Spearman correlation coefficient (color bar) between module z-scores across cells (rows and columns). Module correlations with themselves (diagonals) of 1 were set to “NA” and are shown in grey. **B.** IFN response genes peak early in tumor development. Effect size of differential expression compared to non tumor-bearing lung (color bar, mixed effect logistic regression analysis, **Methods**) for genes (rows) from the IFN response modules 6 and 23 at each timepoint (columns). **C.**Association of T-bet with the IFNstim_TR module 23. Shown is the relation (red curve, loess fit) across cells (dots) between the log_2_(TPM+1) expression (y-axis) of *Tbx21* and the *z*-score of Module 23 (x-axis) in the cell. **D.** T_reg_ module similarity to previously-described expression programs. Spearman correlation coefficient (color bar) between module (columns) *z*-scores across cells and *z*-scores for published signatures (rows, as in **Figure S3A**) of T_reg_ cellular states. **E.**Modules 12 and 21 are enriched for genes of the DP UP signature. Number of genes in the union of modules 12 and 21 (blue), the induced genes in the DP signatures (brown), and their overlap. p < 10^−25^, hypergeometric test. **F.** Example genes whose expression varies significantly over tumor development. Distribution of log_2_(TPM+1) expression of selected genes across time (x-axis). P-value: Kolmogorov-Smirnov test. **G.** DP cells are associated with higher expression of Eff_TR and lower expression of IFNstim_TR genes. ECDF plots of DP vs DN T_reg_ log_2_(fold-change) in gene expression of IFNstim_TR genes (Modules 6 and 23, red, left) or Eff_TR genes (Modules 12 and 21, red, right), and all other genes (gray). P-values: two-sided Kolmogorov – Smirnov tests.

**Figure S5, related to Figure 5.**
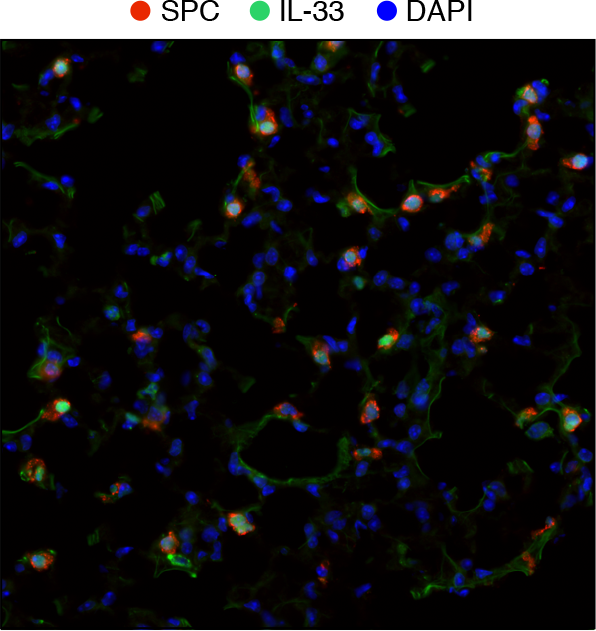
IL-33 is expressed by type II epithelial cells in normal lung. Representative immunofluorescent staining of healthy, non-tumor bearing lung.

**Figure S6, related to Figure 6.**
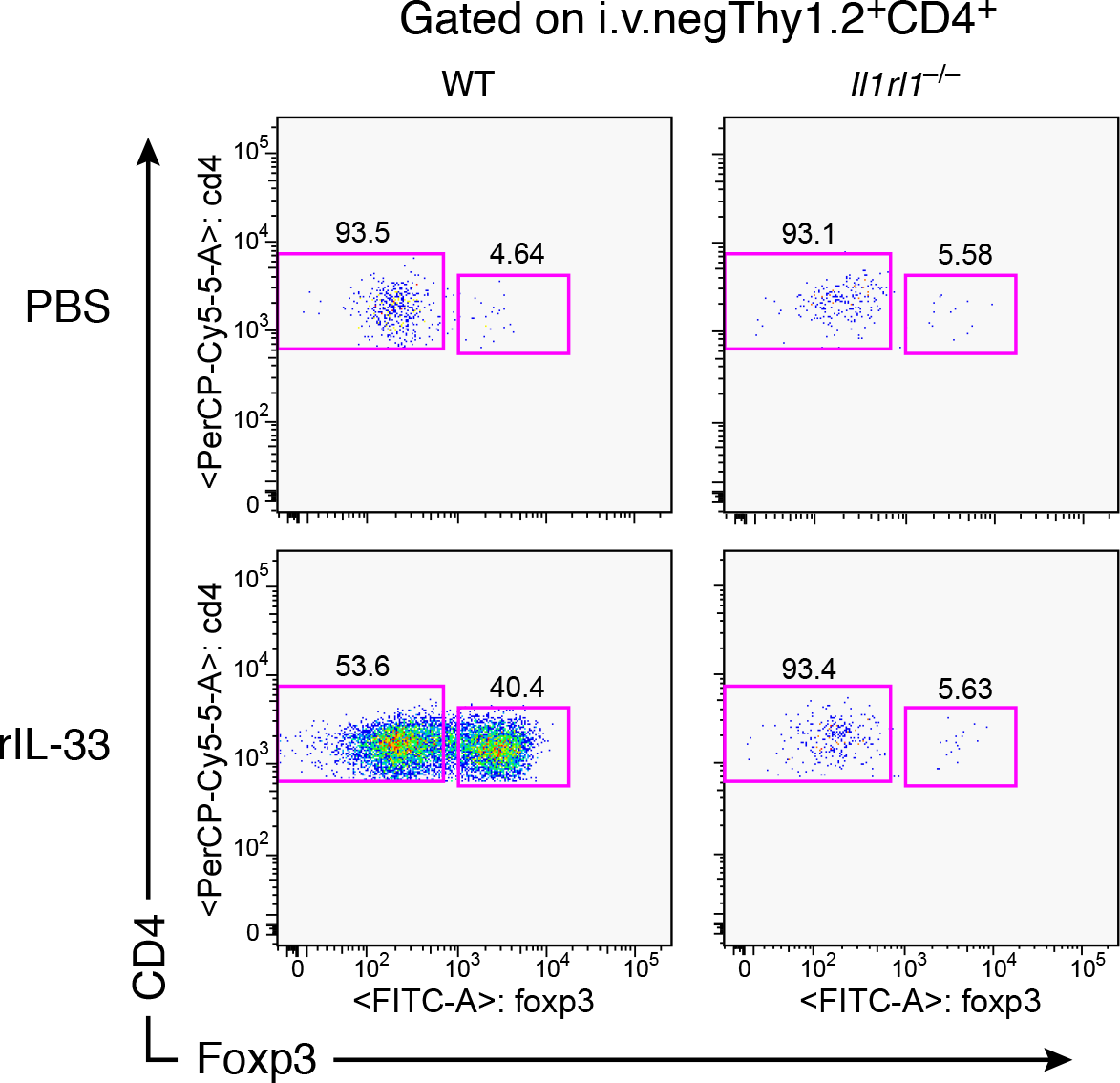
rIL-33 treatment of ST2-deficient mice failed to elicit a change in the proportion of T_regs_. Representative flow cytometry plots of the percentage of T_conv_ (Foxp3^−^) and T_reg_ (Foxp3^+^) of i.v.^neg^Thy1.2^+^CD4^+^ cells in wild-type and ST2-deficient non-tumor bearing mice after challenge with rIL-33 or PBS as control. Data are representative of 2-3 mice per group.

**Figure S7, related to Figure 7.**
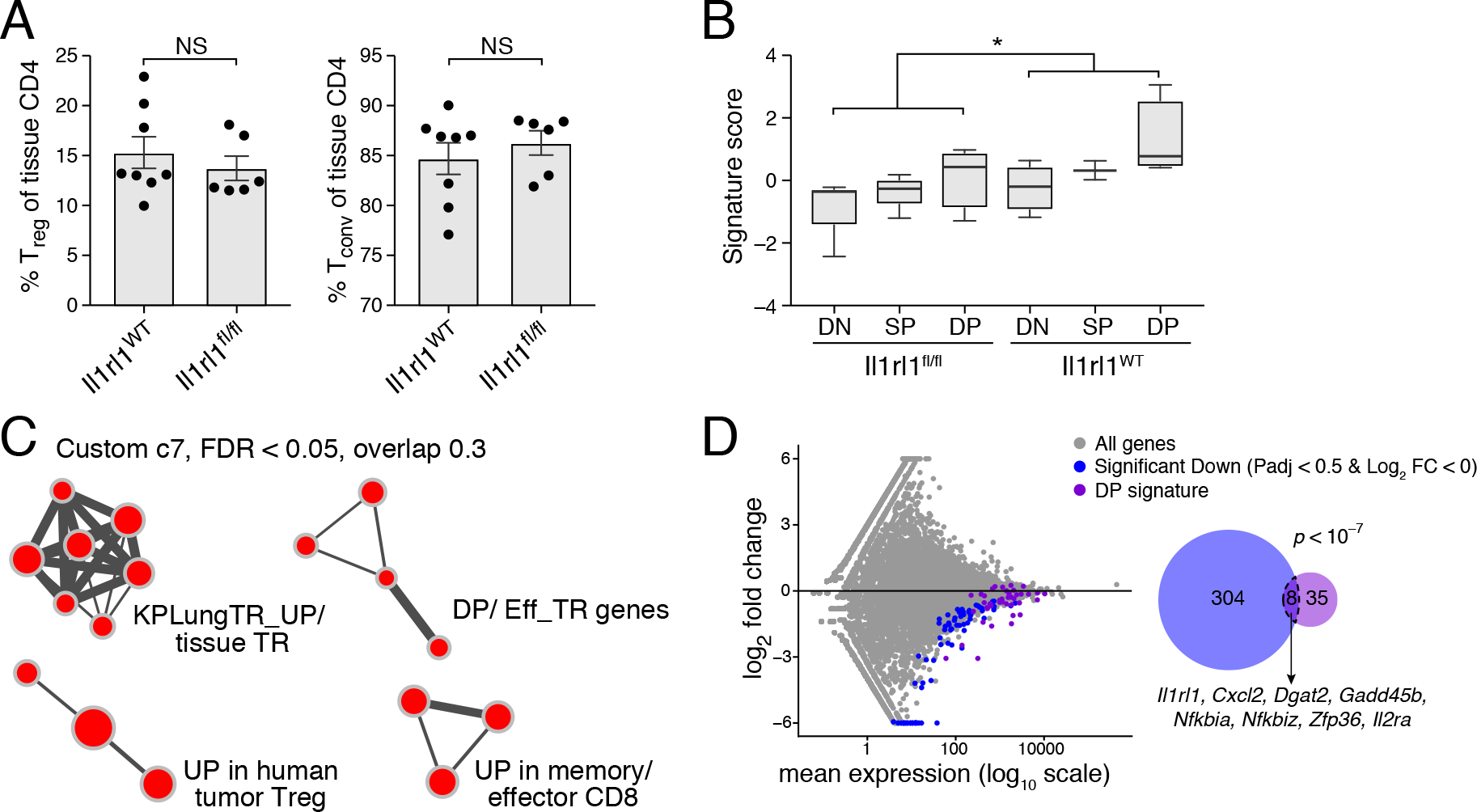
Impact of Treg-specific ST2 ablation on effector T_regs_. **A.** No change in the fraction of T_conv_ or T_regs_ early in tumor development in mice with Treg-specific ST2 deficiency. Percent Foxp3^+^ (left) and %Foxp3^−^(right) of i.v.^neg^CD4^+^ lung cells in KPfrt, *Foxp3*^*YFP-Cre*^ (“*Il1rl1^WT^*”) vs. KPfrt, *Foxp3*^*YFP-Cre*^, *Il1rl1*^*fl/fl*^ (“*Il1rl1^fl/fl^*”) mice at 10 weeks p.i. **B-D.** An expression signature lower in ST2-deficient T_regs_ compared to ST2-wild-type T_regs_ is highest among wild-type DP T_regs_. **B.** Standardized signature score (y-axis) of the expression signature distinguishing *Il1rl1*^*WT*^ and *Il1rl1*^*fl/fl*^ T_regs_ for each lung T_reg_ subpopulation in tumor-bearing mice (x-axis). Box: 25th to 75th percentiles, whiskers: minimum to maximum. Bar: median. No data point is beyond the limit of lines. *p = 0.02, two-sided Mann-Whitney test. **C.** Gene sets enriched in the expression signature distinguishing ST2-deficient T_regs_. GSEA gene sets (nodes) from the custom immune signature database (custom c7, **Methods**) enriched in the signature distinguishing ST2-deficient T_regs_ (p < 0.05, FDR < 0.05; in all significant gene sets. Red: enrichment of upregulated genes. Node size: gene set size. Edge thickness: overlap between gene sets (minimum: 30% overlap). Related pathways were manually annotated. **D.** Left: differential, log_2_(fold change) expression (y-axis) and mean expression (x-axis) for each gene (dot) in CD103^+^KLRG1^+^ (DP) T_regs_ from KPfrt, *Foxp3*^*YFP-Cre*^, *Il1rl1*^*fl/fl*^ vs. KPfrt, *Foxp3*^*YFP-Cre*^ mice. Purple: genes in the DP signature. Blue: Top significantly downregulated genes. Right: Venn diagram shows the overlap between the top differentially downregulated genes in *Il1rl1*^*fl/fl*^ vs. *Il1rl1*^*WT*^ T_regs_ (blue) and the DP signature (purple). P < 10^−7^, hypergeometric test.

## References

Akama-Garren, E.H., Joshi, N.S., Tammela, T., Chang, G.P., Wagner, B.L., Lee, D.-Y., Rideout, W.M. 3rd,, Papagiannakopoulos, T., Xue, W., and Jacks, T. (2016). A Modular Assembly Platform for Rapid Generation of DNA Constructs. Sci. Rep. 6, 16836.

Anderson, K.G., Sung, H., Skon, C.N., Lefrancois, L., Deisinger, A., Vezys, V., and Masopust, D. (2012). Cutting edge: intravascular staining redefines lung CD8 T cell responses. J. Immunol. 189, 2702–2706.

Angerer, P., Haghverdi, L., Büttner, M., Theis, F.J., Marr, C., and Buettner, F. (2016). destiny: diffusion maps for large-scale single-cell data in R. Bioinformatics 32, 1241–1243.

Arpaia, N., Green, J.A., Moltedo, B., Arvey, A., Hemmers, S., Yuan, S., Treuting, P.M., and Rudensky, A.Y. (2015). A Distinct Function of Regulatory T Cells in Tissue Protection. Cell 162, 1078–1089.

Bankhead, P., Loughrey, M.B., Fernández, J.A., Dombrowski, Y., McArt, D.G., Dunne, P.D., McQuaid, S., Gray, R.T., Murray, L.J., Coleman, H.G., et al. (2017). QuPath: Open source software for digital pathology image analysis. Sci. Rep. 7, 16878.

Bettelli, E., Carrier, Y., Gao, W., Korn, T., Strom, T.B., Oukka, M., Weiner, H.L., and Kuchroo, V.K. (2006). Reciprocal developmental pathways for the generation of pathogenic effector TH17 and regulatory T cells. Nature 441, 235–238.

Beyersdorf, N., Ding, X., Tietze, J.K., and Hanke, T. (2007). Characterization of mouse CD4 T cell subsets defined by expression of KLRG1. Eur. J. Immunol. 37, 3445–3454.

Biton, A., Bernard-Pierrot, I., Lou, Y., Krucker, C., Chapeaublanc, E., Rubio-Pérez, C., López-Bigas, N., Kamoun, A., Neuzillet, Y., Gestraud, P., et al. (2014). Independent component analysis uncovers the landscape of the bladder tumor transcriptome and reveals insights into luminal and basal subtypes. Cell Rep. 9, 1235–1245.

Bos, P.D., Plitas, G., Rudra, D., Lee, S.Y., and Rudensky, A.Y. (2013). Transient regulatory T cell ablation deters oncogene-driven breast cancer and enhances radiotherapy. J. Exp. Med. 210, 2435–2466.

Bullard, J.H., Purdom, E., Hansen, K.D., and Dudoit, S. (2010). Evaluation of statistical methods for normalization and differential expression in mRNA-Seq experiments. BMC Bioinformatics 11, 94.

Burzyn, D., Kuswanto, W., Kolodin, D., Shadrach, J.L., Cerletti, M., Jang, Y., Sefik, E., Tan, T.G., Wagers, A.J., Benoist, C., et al. (2013). A special population of regulatory T cells potentiates muscle repair. Cell 155, 1282–1295.

Campbell, D.J. (2015). Control of Regulatory T Cell Migration, Function, and Homeostasis. J. Immunol. 195, 2507–2513.

Caridade, M., Graca, L., and Ribeiro, R.M. (2013). Mechanisms Underlying CD4+ Treg Immune Regulation in the Adult: From Experiments to Models. Front. Immunol. 4, 378.

Cayrol, C., and Girard, J.-P. (2014). IL-33: an alarmin cytokine with crucial roles in innate immunity, inflammation and allergy. Curr. Opin. Immunol. 31, 31–37.

Chen, W.-Y., Hong, J., Gannon, J., Kakkar, R., and Lee, R.T. (2015). Myocardial pressure overload induces systemic inflammation through endothelial cell IL-33. Proc. Natl. Acad. Sci. U. S. A. 112, 7249–7254.

Cheng, G., Yuan, X., Tsai, M.S., Podack, E.R., Yu, A., and Malek, T.R. (2012). IL-2 receptor signaling is essential for the development of Klrg1+ terminally differentiated T regulatory cells. J. Immunol. 189, 1780–1791.

Cipolletta, D., Feuerer, M., Li, A., Kamei, N., Lee, J., Shoelson, S.E., Benoist, C., and Mathis, D. (2012). PPAR-γ is a major driver of the accumulation and phenotype of adipose tissue Treg cells. Nature 486, 549–553.

Der, S.D., Zhou, A., Williams, B.R., and Silverman, R.H. (1998). Identification of genes differentially regulated by interferon alpha, beta, or gamma using oligonucleotide arrays. Proc. Natl. Acad. Sci. U. S. A. 95, 15623–15628.

De Simone, M., Arrigoni, A., Rossetti, G., Gruarin, P., Ranzani, V., Politano, C., Bonnal, R.J.P., Provasi, E., Sarnicola, M.L., Panzeri, I., et al. (2016). Transcriptional Landscape of Human Tissue Lymphocytes Unveils Uniqueness of Tumor-Infiltrating T Regulatory Cells. Immunity 45, 1135–1147.

Dranoff, G. (2011). Experimental mouse tumour models: what can be learnt about human cancer immunology? Nat. Rev. Immunol. 12, 61–66.

DuPage, M., Dooley, A.L., and Jacks, T. (2009). Conditional mouse lung cancer models using adenoviral or lentiviral delivery of Cre recombinase. Nat. Protoc. 4, 1064–1072.

DuPage, M., Cheung, A.F., Mazumdar, C., Winslow, M.M., Bronson, R., Schmidt, L.M., Crowley, D., Chen, J., and Jacks, T. (2011). Endogenous T cell responses to antigens expressed in lung adenocarcinomas delay malignant tumor progression. Cancer Cell 19, 72–85.

Engel, I., Seumois, G., Chavez, L., Samaniego-Castruita, D., White, B., Chawla, A., Mock, D., Vijayanand, P., and Kronenberg, M. (2016). Innate-like functions of natural killer T cell subsets result from highly divergent gene programs. Nat. Immunol. 17, 728–739.

Fan, J., Salathia, N., Liu, R., Kaeser, G.E., Yung, Y.C., Herman, J.L., Kaper, F., Fan, J.-B., Zhang, K., Chun, J., et al. (2016). Characterizing transcriptional heterogeneity through pathway and gene set overdispersion analysis. Nat. Methods 13, 241–244.

Feuerer, M., Herrero, L., Cipolletta, D., Naaz, A., Wong, J., Nayer, A., Lee, J., Goldfine, A.B., Benoist, C., Shoelson, S., et al. (2009). Lean, but not obese, fat is enriched for a unique population of regulatory T cells that affect metabolic parameters. Nat. Med. 15, 930–939.

Fridman, W.H., Pagès, F., Sautès-Fridman, C., and Galon, J. (2012). The immune contexture in human tumours: impact on clinical outcome. Nat. Rev. Cancer 12, 298–306.

Gibson, D.G., Young, L., Chuang, R.-Y., Venter, J.C., Hutchison, C.A. 3rd,, and Smith, H.O. (2009). Enzymatic assembly of DNA molecules up to several hundred kilobases. Nat. Methods 6, 343–345.

Green, J.A., Arpaia, N., Schizas, M., Dobrin, A., and Rudensky, A.Y. (2017). A nonimmune function of T cells in promoting lung tumor progression. J. Exp. Med. 214, 3565–3575.

Guo, X., Zhang, Y., Zheng, L., Zheng, C., Song, J., Zhang, Q., Kang, B., Liu, Z., Jin, L., Xing, R., et al. (2018). Global characterization of T cells in non-small-cell lung cancer by single-cell sequencing. Nat. Med. 24, 978–985.

Halim, L., Romano, M., McGregor, R., Correa, I., Pavlidis, P., Grageda, N., Hoong, S.-J., Yuksel, M., Jassem, W., Hannen, R.F., et al. (2017). An Atlas of Human Regulatory T Helper-like Cells Reveals Features of Th2-like Tregs that Support a Tumorigenic Environment. Cell Rep. 20, 757–770.

Hall, A.O., Beiting, D.P., Tato, C., John, B., Oldenhove, G., Lombana, C.G., Pritchard, G.H., Silver, J.S., Bouladoux, N., Stumhofer, J.S., et al. (2012). The cytokines interleukin 27 and interferon-γ promote distinct Treg cell populations required to limit infection-induced pathology. Immunity 37, 511–523.

Huehn, J., Siegmund, K., Lehmann, J.C.U., Siewert, C., Haubold, U., Feuerer, M., Debes, G.F., Lauber, J., Frey, O., Przybylski, G.K., et al. (2004). Developmental stage, phenotype, and migration distinguish naive- and effector/memory-like CD4+ regulatory T cells. J. Exp. Med. 199, 303–313.

Hyvärinen, A., and Oja, E. (2000). Independent component analysis: algorithms and applications. Neural Netw. 13, 411–430.

Jackson, E.L., Olive, K.P., Tuveson, D.A., Bronson, R., Crowley, D., Brown, M., and Jacks, T. (2005). The differential effects of mutant p53 alleles on advanced murine lung cancer. Cancer Res. 65, 10280–10288.

Johdi, N.A., Ait-Tahar, K., Sagap, I., and Jamal, R. (2017). Molecular Signatures of Human Regulatory T Cells in Colorectal Cancer and Polyps. Front. Immunol. 8, 620.

Josefowicz, S.Z., Lu, L.-F., and Rudensky, A.Y. (2012). Regulatory T cells: mechanisms of differentiation and function. Annu. Rev. Immunol. 30, 531–564.

Joshi, N.S., Akama-Garren, E.H., Lu, Y., Lee, D.-Y., Chang, G.P., Li, A., DuPage, M., Tammela, T., Kerper, N.R., Farago, A.F., et al. (2015). Regulatory T Cells in Tumor-Associated Tertiary Lymphoid Structures Suppress Anti-tumor T Cell Responses. Immunity 43, 579–590.

Jovanovic, I.P., Pejnovic, N.N., Radosavljevic, G.D., Pantic, J.M., Milovanovic, M.Z., Arsenijevic, N.N., and Lukic, M.L. (2014). Interleukin-33/ST2 axis promotes breast cancer growth and metastases by facilitating intratumoral accumulation of immunosuppressive and innate lymphoid cells. Int. J. Cancer 134, 1669–1682.

Kim, B.-S., Lu, H., Ichiyama, K., Chen, X., Zhang, Y.-B., Mistry, N.A., Tanaka, K., Lee, Y.-H., Nurieva, R., Zhang, L., et al. (2017). Generation of RORγt+ Antigen-Specific T Regulatory 17 Cells from Foxp3+ Precursors in Autoimmunity. Cell Rep. 21, 195–207.

Kim, J.M., Rasmussen, J.P., and Rudensky, A.Y. (2007). Regulatory T cells prevent catastrophic autoimmunity throughout the lifespan of mice. Nat. Immunol. 8, 191–197.

Koch, M.A., Tucker-Heard, G.’s, Perdue, N.R., Killebrew, J.R., Urdahl, K.B., and Campbell, D.J. (2009). The transcription factor T-bet controls regulatory T cell homeostasis and function during type 1 inflammation. Nat. Immunol. 10, 595–602.

Koch, M.A., Thomas, K.R., Perdue, N.R., Smigiel, K.S., Srivastava, S., and Campbell, D.J. (2012). T-bet(+) Treg cells undergo abortive Th1 cell differentiation due to impaired expression of IL-12 receptor β2. Immunity 37, 501–510.

Kohlmeier, J.E., Cookenham, T., Miller, S.C., Roberts, A.D., Christensen, J.P., Thomsen, A.R., and Woodland, D.L. (2009). CXCR3 directs antigen-specific effector CD4+ T cell migration to the lung during parainfluenza virus infection. J. Immunol. 183, 4378–4384.

Kolodin, D., van Panhuys, N., Li, C., Magnuson, A.M., Cipolletta, D., Miller, C.M., Wagers, A., Germain, R.N., Benoist, C., and Mathis, D. (2015). Antigen- and cytokine-driven accumulation of regulatory T cells in visceral adipose tissue of lean mice. Cell Metab. 21, 543–557.

Kondo, Y., Yoshimoto, T., Yasuda, K., Futatsugi-Yumikura, S., Morimoto, M., Hayashi, N., Hoshino, T., Fujimoto, J., and Nakanishi, K. (2008). Administration of IL-33 induces airway hyperresponsiveness and goblet cell hyperplasia in the lungs in the absence of adaptive immune system. Int. Immunol. 20, 791–800.

Kuswanto, W., Burzyn, D., Panduro, M., Wang, K.K., Jang, Y.C., Wagers, A.J., Benoist, C., and Mathis, (2016). Poor Repair of Skeletal Muscle in Aging Mice Reflects a Defect in Local, Interleukin-33-Dependent Accumulation of Regulatory T Cells. Immunity 44, 355–367.

Langmead, B., Trapnell, C., Pop, M., and Salzberg, S.L. (2009). Ultrafast and memory-efficient alignment of short DNA sequences to the human genome. Genome Biol. 10, R25.

Lee, Y., Awasthi, A., Yosef, N., Quintana, F.J., Xiao, S., Peters, A., Wu, C., Kleinewietfeld, M., Kunder, S., Hafler, D.A., et al. (2012). Induction and molecular signature of pathogenic TH17 cells. Nat. Immunol. 13, 991–999.

Lee, Y.K., Turner, H., Maynard, C.L., Oliver, J.R., Chen, D., Elson, C.O., and Weaver, C.T. (2009). Late developmental plasticity in the T helper 17 lineage. Immunity 30, 92–107.

Lehmann, J., Huehn, J., de la Rosa, M., Maszyna, F., Kretschmer, U., Krenn, V., Brunner, M., Scheffold, A., and Hamann, A. (2002). Expression of the integrin alpha Ebeta 7 identifies unique subsets of CD25+ as well as CD25-regulatory T cells. Proc. Natl. Acad. Sci. U. S. A. 99, 13031–13036.

Levine, A.G., Mendoza, A., Hemmers, S., Moltedo, B., Niec, R.E., Schizas, M., Hoyos, B.E., Putintseva, V., Chaudhry, A., Dikiy, S., et al. (2017). Stability and function of regulatory T cells expressing the transcription factor T-bet. Nature 546, 421–425.

Li, B., and Dewey, C.N. (2011). RSEM: accurate transcript quantification from RNA-Seq data with or without a reference genome. BMC Bioinformatics 12, 323.

Li, M.O., and Rudensky, A.Y. (2016). T cell receptor signalling in the control of regulatory T cell differentiation and function. Nat. Rev. Immunol. 16, 220–233.

Li, D., Guabiraba, R., Besnard, A.-G., Komai-Koma, M., Jabir, M.S., Zhang, L., Graham, G.J., Kurowska-Stolarska, M., Liew, F.Y., McSharry, C., et al. (2014). IL-33 promotes ST2-dependent lung fibrosis by the induction of alternatively activated macrophages and innate lymphoid cells in mice. J. Allergy Clin. Immunol. 134, 1422–1432.e11.

Li, L., Zeng, Q., Bhutkar, A., Galván, J.A., Karamitopoulou, E., Noordermeer, D., Peng, M.-W., Piersigilli, A., Perren, A., Zlobec, I., et al. (2018). GKAP Acts as a Genetic Modulator of NMDAR Signaling to Govern Invasive Tumor Growth. Cancer Cell 33, 736–751.e5.

Magnuson, A.M., Kiner, E., Ergun, A., Park, J.S., Asinovski, N., Ortiz-Lopez, A., Kilcoyne, A., Paoluzzi-Tomada, E., Weissleder, R., Mathis, D., et al. (2018). Identification and validation of a tumor-infiltrating Treg transcriptional signature conserved across species and tumor types. Proc. Natl. Acad. Sci. U. S. A.

Makkouk, A., and Weiner, G.J. (2015). Cancer immunotherapy and breaking immune tolerance: new approaches to an old challenge. Cancer Res. 75, 5–10.

Marabelle, A., Kohrt, H., Sagiv-Barfi, I., Ajami, B., Axtell, R.C., Zhou, G., Rajapaksa, R., Green, M.R., Torchia, J., Brody, J., et al. (2013). Depleting tumor-specific Tregs at a single site eradicates disseminated tumors. J. Clin. Invest. 123, 2447–2463.

Miragaia, R.J., Gomes, T., Chomka, A., Jardine, L., Riedel, A., Hegazy, A.N., Lindeman, I., Emerton, G., Krausgruber, T., Shields, J., et al. (2017). Single cell transcriptomics of regulatory T cells reveals trajectories of tissue adaptation. bioRxiv 217489.

Nordhausen, K., Cardoso, J.F., Miettinen, J., Oja, H., Ollila, E., and Taskinen, S. (2014). JADE: JADE and other BSS methods as well as some BSS performance criteria. R Package Version 1–9.

Overacre-Delgoffe, A.E., Chikina, M., Dadey, R.E., Yano, H., Brunazzi, E.A., Shayan, G., Horne, W., Moskovitz, J.M., Kolls, J.K., Sander, C., et al. (2017). Interferon-γ Drives Treg Fragility to Promote Anti-tumor Immunity. Cell 169, 1130–1141.e11.

Petersen, R.P., Campa, M.J., Sperlazza, J., Conlon, D., Joshi, M.-B., Harpole, D.H. Jr,, and Patz, E.F. Jr, (2006). Tumor infiltrating Foxp3+ regulatory T-cells are associated with recurrence in pathologic stage I NSCLC patients. Cancer 107, 2866–2872.

Picelli, S., Björklund, Å.K., Faridani, O.R., Sagasser, S., Winberg, G., and Sandberg, R. (2013). Smart-seq2 for sensitive full-length transcriptome profiling in single cells. Nat. Methods 10, 1096–1098.

Plitas, G., Konopacki, C., Wu, K., Bos, P.D., Morrow, M., Putintseva, E.V., Chudakov, D.M., and Rudensky, A.Y. (2016). Regulatory T Cells Exhibit Distinct Features in Human Breast Cancer. Immunity 45, 1122–1134.

Redjimi, N., Raffin, C., Raimbaud, I., Pignon, P., Matsuzaki, J., Odunsi, K., Valmori, D., and Ayyoub, M. (2012). CXCR3+ T regulatory cells selectively accumulate in human ovarian carcinomas to limit type I immunity. Cancer Res. 72, 4351–4360.

Rutledge, D.N., and Jouan-Rimbaud Bouveresse, D. (2013). Independent Components Analysis with the JADE algorithm. Trends Analyt. Chem. 50, 22–32.

Saito, T., Nishikawa, H., Wada, H., Nagano, Y., Sugiyama, D., Atarashi, K., Maeda, Y., Hamaguchi, M., Ohkura, N., Sato, E., et al. (2016). Two FOXP3(+)CD4(+) T cell subpopulations distinctly control the prognosis of colorectal cancers. Nat. Med. 22, 679–684.

Sakaguchi, S. (2011). Regulatory T cells: history and perspective. Methods Mol. Biol. 707, 3–17.

Sánchez-Rivera, F.J., Papagiannakopoulos, T., Romero, R., Tammela, T., Bauer, M.R., Bhutkar, A., Joshi, N.S., Subbaraj, L., Bronson, R.T., Xue, W., et al. (2014). Rapid modelling of cooperating genetic events in cancer through somatic genome editing. Nature 516, 428–431.

Sather, B.D., Treuting, P., Perdue, N., Miazgowicz, M., Fontenot, J.D., Rudensky, A.Y., and Campbell, D.J. (2007). Altering the distribution of Foxp3(+) regulatory T cells results in tissue-specific inflammatory disease. J. Exp. Med. 204, 1335–1347.

Savage, P.A., Malchow, S., and Leventhal, D.S. (2013). Basic principles of tumor-associated regulatory T cell biology. Trends Immunol. 34, 33–40.

Schiering, C., Krausgruber, T., Chomka, A., Fröhlich, A., Adelmann, K., Wohlfert, E.A., Pott, J., Griseri, T., Bollrath, J., Hegazy, A.N., et al. (2014). The alarmin IL-33 promotes regulatory T-cell function in the intestine. Nature 513, 564–568.

Schmitz, J., Owyang, A., Oldham, E., Song, Y., Murphy, E., McClanahan, T.K., Zurawski, G., Moshrefi, M., Qin, J., Li, X., et al. (2005). IL-33, an interleukin-1-like cytokine that signals via the IL-1 receptor-related protein ST2 and induces T helper type 2-associated cytokines. Immunity 23, 479–490.

Shalek, A.K., Satija, R., Adiconis, X., Gertner, R.S., Gaublomme, J.T., Raychowdhury, R., Schwartz, S., Yosef, N., Malboeuf, C., Lu, D., et al. (2013). Single-cell transcriptomics reveals bimodality in expression and splicing in immune cells. Nature 498, 236–240.

Shang, B., Liu, Y., Jiang, S.-J., and Liu, Y. (2015). Prognostic value of tumor-infiltrating FoxP3^+^ regulatory T cells in cancers: a systematic review and meta-analysis. Sci. Rep. 5, 15179.

Shimizu, K., Nakata, M., Hirami, Y., Yukawa, T., Maeda, A., and Tanemoto, K. (2010). Tumor-infiltrating Foxp3+ regulatory T cells are correlated with cyclooxygenase-2 expression and are associated with recurrence in resected non-small cell lung cancer. J. Thorac. Oncol. 5, 585–590.

Shugay, M., Bagaev, D.V., Turchaninova, M.A., Bolotin, D.A., Britanova, O.V., Putintseva, E.V., Pogorelyy, M.V., Nazarov, V.I., Zvyagin, I.V., Kirgizova, V.I., et al. (2015). VDJtools: Unifying Post-analysis of T Cell Receptor Repertoires. PLoS Comput. Biol. 11, e1004503.

Simpson, T.R., Li, F., Montalvo-Ortiz, W., Sepulveda, M.A., Bergerhoff, K., Arce, F., Roddie, C., Henry, J.Y., Yagita, H., Wolchok, J.D., et al. (2013). Fc-dependent depletion of tumor-infiltrating regulatory T cells co-defines the efficacy of anti-CTLA-4 therapy against melanoma. J. Exp. Med. 210, 1695–1710.

Singer, M., Wang, C., Cong, L., Marjanovic, N.D., Kowalczyk, M.S., Zhang, H., Nyman, J., Sakuishi, K., Kurtulus, S., Gennert, D., et al. (2017). A Distinct Gene Module for Dysfunction Uncoupled from Activation in Tumor-Infiltrating T Cells. Cell 171, 1221–1223.

Soria, J.-C., Marabelle, A., Brahmer, J.R., and Gettinger, S. (2015). Immune Checkpoint Modulation for Non-Small Cell Lung Cancer. Clin. Cancer Res. 21, 2256–2262.

Stubbington, M.J.T., Lönnberg, T., Proserpio, V., Clare, S., Speak, A.O., Dougan, G., and Teichmann, S.A. (2016). T cell fate and clonality inference from single-cell transcriptomes. Nat. Methods 13, 329–332.

Subramanian, A., Tamayo, P., Mootha, V.K., Mukherjee, S., Ebert, B.L., Gillette, M.A., Paulovich, A., Pomeroy, S.L., Golub, T.R., Lander, E.S., et al. (2005). Gene set enrichment analysis: a knowledge-based approach for interpreting genome-wide expression profiles. Proc. Natl. Acad. Sci. U. S. A. 102, 15545–15550.

Suzuki, K., Kadota, K., Sima, C.S., Nitadori, J.-I., Rusch, V.W., Travis, W.D., Sadelain, M., and Adusumilli, P.S. (2013). Clinical impact of immune microenvironment in stage I lung adenocarcinoma: tumor interleukin-12 receptor β2 (IL-12Rβ2), IL-7R, and stromal FoxP3/CD3 ratio are independent predictors of recurrence. J. Clin. Oncol. 31, 490–498.

Tanaka, A., and Sakaguchi, S. (2017). Regulatory T cells in cancer immunotherapy. Cell Res. 27, 109–118.

Townsend, M.J., Fallon, P.G., Matthews, D.J., Jolin, H.E., and McKenzie, A.N. (2000). T1/ST2-deficient mice demonstrate the importance of T1/ST2 in developing primary T helper cell type 2 responses. J. Exp. Med. 191, 1069–1076.

Vasanthakumar, A., Moro, K., Xin, A., Liao, Y., Gloury, R., Kawamoto, S., Fagarasan, S., Mielke, L.A., Afshar-Sterle, S., Masters, S.L., et al. (2015). The transcriptional regulators IRF4, BATF and IL-33 orchestrate development and maintenance of adipose tissue-resident regulatory T cells. Nat. Immunol. 16, 276–285.

van der Veeken, J., Gonzalez, A.J., Cho, H., Arvey, A., Hemmers, S., Leslie, C.S., and Rudensky, A.Y. (2016). Memory of Inflammation in Regulatory T Cells. Cell 166, 977–990.

Végran, F., Apetoh, L., and Ghiringhelli, F. (2015). Th9 cells: a novel CD4 T-cell subset in the immune war against cancer. Cancer Res. 75, 475–479.

Vignali, D.A.A., Collison, L.W., and Workman, C.J. (2008). How regulatory T cells work. Nat. Rev. Immunol. 8, 523–532.

Walker, J.A., and McKenzie, A.N.J. (2018). TH2 cell development and function. Nat. Rev. Immunol. 18, 121–133.

Wan, Y.Y., and Flavell, R.A. (2005). Identifying Foxp3-expressing suppressor T cells with a bicistronic reporter. Proc. Natl. Acad. Sci. U. S. A. 102, 5126–5131.

Wang, C., Chen, Z., Bu, X., Han, Y., Shan, S., Ren, T., and Song, W. (2016). IL-33 signaling fuels outgrowth and metastasis of human lung cancer. Biochem. Biophys. Res. Commun. 479, 461–468.

Wang, K., Shan, S., Yang, Z., Gu, X., Wang, Y., Wang, C., and Ren, T. (2017). IL-33 blockade suppresses tumor growth of human lung cancer through direct and indirect pathways in a preclinical model. Oncotarget 8, 68571–68582.

Wang, Y., Godec, J., Ben-Aissa, K., Cui, K., Zhao, K., Pucsek, A.B., Lee, Y.K., Weaver, C.T., Yagi, R., and Lazarevic, V. (2014). The transcription factors T-bet and Runx are required for the ontogeny of pathogenic interferon-γ-producing T helper 17 cells. Immunity 40, 355–366.

Yamazaki, T., Yang, X.O., Chung, Y., Fukunaga, A., Nurieva, R., Pappu, B., Martin-Orozco, N., Kang, H.S., Ma, L., Panopoulos, A.D., et al. (2008). CCR6 regulates the migration of inflammatory and regulatory T cells. J. Immunol. 181, 8391–8401.

Young, N.P., Crowley, D., and Jacks, T. (2011). Uncoupling cancer mutations reveals critical timing of p53 loss in sarcomagenesis. Cancer Res. 71, 4040–4047.

Zheng, C., Zheng, L., Yoo, J.-K., Guo, H., Zhang, Y., Guo, X., Kang, B., Hu, R., Huang, J.Y., Zhang, Q., et al. (2017). Landscape of Infiltrating T Cells in Liver Cancer Revealed by Single-Cell Sequencing. Cell 169, 1342–1356.e16.

